# Widespread sex-dimorphism across single-cell transcriptomes of adult African turquoise killifish tissues

**DOI:** 10.1101/2023.05.05.539616

**Authors:** Bryan B. Teefy, Aaron J.J. Lemus, Ari Adler, Alan Xu, Rajyk Bhala, Katelyn Hsu, Bérénice A. Benayoun

**Affiliations:** Leonard Davis School of Gerontology, University of Southern California, Los Angeles, CA 90089, USA; Molecular and Computational Biology Department, USC Dornsife College of Letters, Arts and Sciences, Los Angeles, CA 90089, USA; Quantitative & Computational Biology Department, USC Dornsife College of Letters, Arts and Sciences, Los Angeles, CA 90089, USA; Biochemistry and Molecular Medicine Department, USC Keck School of Medicine, Los Angeles, CA 90089, USA; USC Norris Comprehensive Cancer Center, Epigenetics and Gene Regulation, Los Angeles, CA 90089, USA; USC Stem Cell Initiative, Los Angeles, CA 90089, USA

## Abstract

The African turquoise killifish (*Nothobranchius furzeri*), the shortest-lived vertebrate that can be bred in captivity, is an emerging model organism to study vertebrate aging. Here we describe the first multi-tissue, single-cell gene expression atlas of female and male turquoise killifish tissues comprising immune and metabolic cells from the blood, kidney, liver, and spleen. We were able to annotate 22 distinct cell types, define associated marker genes, and infer differentiation trajectories. Using this dataset, we found pervasive sex-dimorphic gene expression across cell types, especially in the liver. Sex-dimorphic genes tended to be involved in processes related to lipid metabolism, and indeed, we observed clear differences in lipid storage in female *vs*. male turquoise killifish livers. Importantly, we use machine-learning to predict sex using single-cell gene expression in our atlas and identify potential transcriptional markers for molecular sex identity in this species. As proof-of-principle, we show that our atlas can be used to deconvolute existing liver bulk RNA-seq data in this species to obtain accurate estimates of cell type proportions across biological conditions. We believe that this single-cell atlas can be a resource to the community that could notably be leveraged to identify cell type-specific genes for cell type-specific expression in transgenic animals.

## Introduction

Aging is a multi-factorial decline in fitness that is the major risk factor for a vast array of diseases^1^. One model uniquely positioned to accelerate research into the drivers of organismal aging is the African turquoise killifish (*Nothobranchius furzeri*). The turquoise killifish is the shortest-lived vertebrate that can be bred in captivity, which makes longitudinal studies in vertebrates more time- and cost-effective than other more traditional vertebrate models^2, 3^. Furthermore, the turquoise killifish is one of a small number of non-traditional model organisms that is amenable to transgenesis^2–9^. Though molecular biology is now advancing rapidly in this species, a definitive characterization of turquoise killifish cell types, especially those within tissues that are known to be impacted drastically by the aging process, remains to be established.

Aging is characterized by a set of conserved “hallmarks”, including (i) chronic inflammation and (ii) deregulated nutrient sensing^10^. Indeed, response to pathogen exposure becomes defective with increasing age across species^11^. This is concomitant with an increased baseline expression in innate immunity-related genes even in the absence of overt infection, a phenomenon dubbed “sterile inflammation” or “inflammaging”^12^. Deregulated nutrient sensing (including through altered insulin signaling) is common in aged animals and promotes a generalized metabolic decline^13^. Interventions that decrease growth and metabolic rates (*e*.*g.* caloric restriction) increase healthspan and lifespan across species^14^, including in the turquoise killifish ^15^. Interestingly, both immune responses^16^ and metabolism^17^ are highly influenced by sex, although such differences are still not widely studied, especially in large ‘omic’ profiling efforts.

In teleost fish, hematopoiesis- and therefore immune cell biogenesis-largely occurs in the kidney. Within the kidney, hematopoiesis occurs in a substructure known as the head kidney, which is considered the functional equivalent of the mammalian bone marrow^18–20^. In some teleost species, the spleen is also involved in hematopoiesis, typically as the primary erythropoietic organ^21, 22^. Thus, the kidney, spleen, and blood are rich targets from which to explore immune biology in the turquoise killifish. Regarding metabolism, the liver is the predominant metabolic organ in vertebrates, including teleosts^23, 24^. Since metabolism is known to change drastically with age, the liver is an optimal organ for studying metabolic aging ^25^.

In this study, we generated a multi-tissue single-cell gene expression atlas across key immune and metabolic tissues in the African turquoise killifish. For this study, we focused on profiling single cells derived from blood, kidney, liver, and spleen of adult female and male fish. Using a combination of automated annotation using markers from other vertebrate species and manual curation, we defined distinct cell types and propose robust specific marker genes for each cell type. Since our atlas has equal representation from both sexes, we interrogate cell type specific sex-dimorphic gene expression signatures, revealing a predominance of genes involved in lipid metabolism. We show that we can predict sex from gene expression data, providing a potential marker gene for quality control of future transcriptomic studies in this species. Finally, we provide a proof-of-principle analysis that our atlas can be used to accurately deconvolute existing RNA-seq datasets into their component cell types. This transcriptomic atlas will serve as a resource for the study of immunity, metabolism, and sex-dimorphism in the African turquoise killifish.

## Results

### A single-cell atlas of gene expression across African turquoise killifish tissues

To annotate cell types and understand cell type specific gene expression in this species, we performed single-cell RNA sequencing on cells isolated from blood, kidney, liver, and spleens from 6-week-old male and female African turquoise killifish from the GRZ strain (**Figure 1A**). In our husbandry conditions, this age corresponds to sexually mature adults, generally at peak health^26^. Relevant solid tissues were harvested and pooled from 3-4 fish of the same sex to generate enough material for sex-specific single-cell sequencing. We optimized a dissociation protocol that consistently isolated cells from solid tissues with a high singlet rate (>95%) as measured by flow cytometry (see Methods; **Supplementary Figure S1A, Supplementary Table S1A**). We isolated cells from 3 independent cohorts of turquoise killifish, for which cell counts and viability were measured with a cell counter using trypan blue staining (**Supplementary Table S1B**). Only samples with sufficient viability were included for downstream processing for droplet microfluidics-based single-cell RNA-seq profiling using the 10xGenomics platform (**Supplementary Figure S1B, Supplementary Table S1B**). After standard pre-processing, we detected 112,155 cells, achieving a mean sequencing depth of ∼45,435 reads per cell at a median sequence saturation rate of 62.3% across libraries (**Supplementary Table S2**). Importantly, since the turquoise killifish genome is rich in transposable elements (TEs) ^26–28^, we analyzed single-cell transcriptomes for both gene and TE-derived transcripts (see **Methods**).

**Figure 1.**
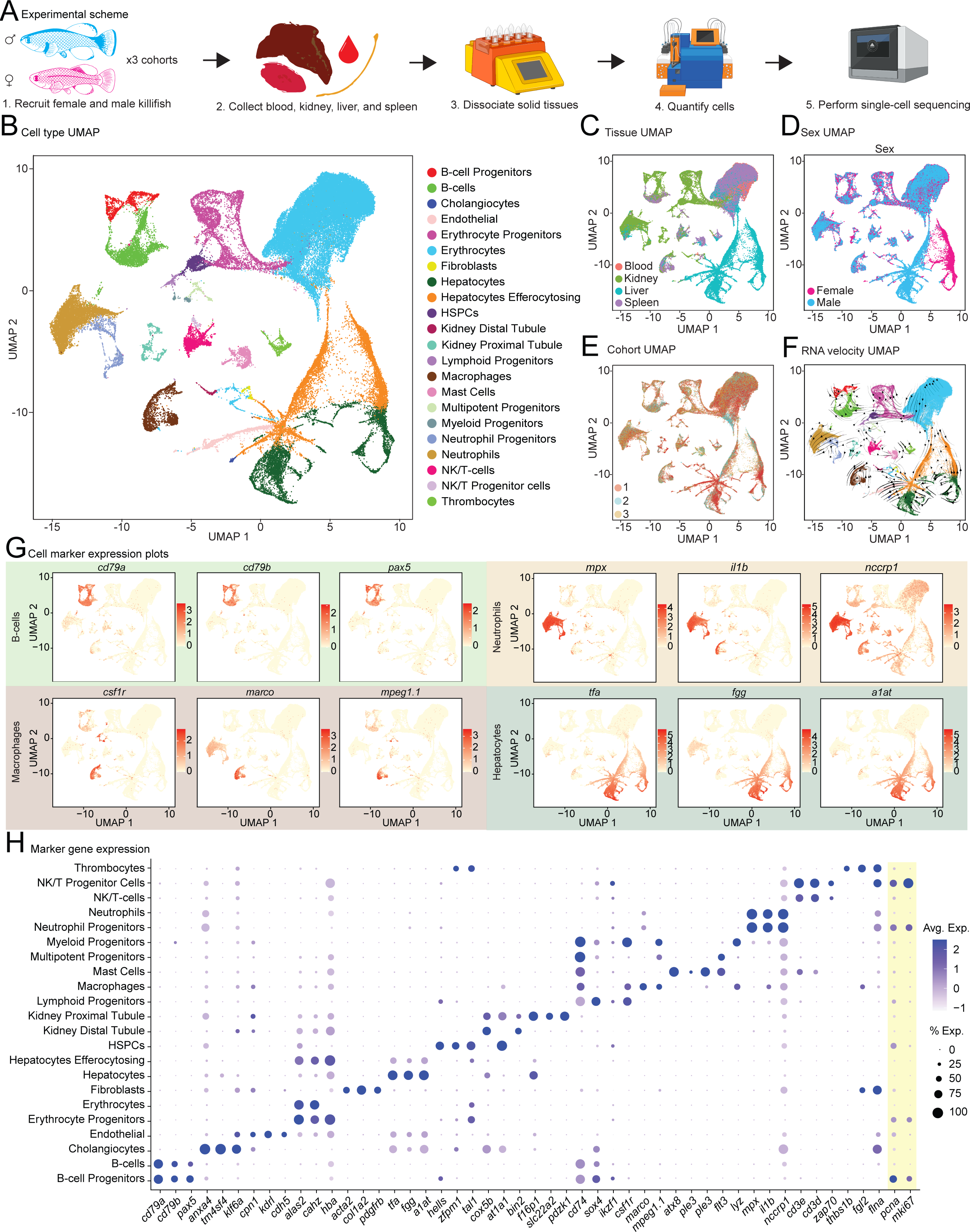
A single-cell atlas of female and male African turquoise killifish tissues. **(A)** Schematic for tissue dissociation and sequencing. The blood, kidneys, livers, and spleens were isolated from 3 cohorts of 3-4 fish and solid tissues were dissociated into single-cell suspensions. Single-cell RNA-seq libraries were made using a standard 10xGenomics protocol. **(B-F)** UMAP plots of the killifish tissue atlas denoted by **(B)** annotated cell type, **(C)** tissue of origin, **(D)** sex, **(E)** cohort, and **(F)** RNA velocity. **(G)** Expression UMAP for canonical marker genes for B-cells, macrophages, neutrophils, and hepatocytes. **(H)** Dotplot of marker gene expression, showing the average expression level and percentage of cells expressing each marker gene. Markers of cell proliferation (*pcna* and *mki67*) are provided under a yellow overlay, since they were used to annotated proliferative stem/progenitor states. Since many NCBI gene names were uninformative, names corresponding to their predicted homologs are used. Original NCBI gene names are provided in **Supplementary Table S9**. HSPCs: hematopoietic stem and progenitor cells.

After quality filtering and doublet removal, we retained 81,357 high quality cells across our 3 experimental cohorts (see **Methods**). Using unsupervised clustering together with markers derived from Zebrafish cell atlases^29–37^ (**Supplementary Table S3**), we identified 22 distinct cell types, including key stem and mature cell types across tissues and sexes across all 3 cohorts (**Figure 1B-E**). Cell types were annotated primarily using automated cell annotation software, with remaining ambiguous clusters annotated after manual inspection (see **Methods**). Importantly, annotated cell types showed distinct expression of specific canonical cell type markers (**Figure 1G, H,** and **Supplementary Figure S1D**).

We selected tissues for profiling in our atlas in part to enable the capture of hematopoiesis, which primarily occurs in the kidney in teleosts^18–20^. Thus, to infer differentiation trajectories, we performed RNA velocity analysis^38^ on our atlas to annotate potential trajectories in the killifish hematopoietic system (**Figure 1F**). Hematopoietic stem and progenitor cells (HSPCs) do not have an apparent “velocity” and can be observed differentiating into myeloid and lymphoid cell clusters. Neutrophils, which develop rapidly from their progenitor cells, have large velocity vectors emanating from the neutrophil progenitors towards a presumed mature neutrophil state^39^. Unexpectedly, erythrocyte progenitors appear to be differentiating into HSPCs. However, this is a known artifact of RNA velocity analysis^40^. Indeed, erythrocyte differentiation begins with a rapid burst of transcription, which appears as “reverse” development to the RNA velocity algorithm^40^.

Although every other cell cluster contained cells from animals for both sexes, we noted that cells from female *vs*. male turquoise killifish livers occupied different spaces in the UMAP space (**Figure 1C, D**). This pattern held for each independent cohort (**Figure 1E**), making it unlikely to represent a technical artifact. The liver tissue of female teleosts has been shown to produce yolk biomolecules necessary for successful oogenesis, *e.g.* the yolk precursor glycolipophosphoprotein vitellogenin^41, 42^. Consistently, in our dataset, female hepatocytes show high expression of *vitellogenin-1-like*, unlike their male counterpart (**Supplementary Figure S1C**). Although other cell types, including cholangiocytes and endothelial cells, also express lower levels of *vitellogenin-1-like*, expression of this gene is strictly restricted to the liver (**Supplementary Figure S1C**). Intriguingly, a major liver cell type robustly expressed markers of both hepatocytes and erythrocytes across all 3 cohorts and both sexes (**Figure 1H** and **Supplementary Figure S1D**). Since hepatocytes can perform efferocytosis of dead/dying cells^43^, we hypotehsize that these cells are likely hepatocytes engulfing dead/dying erythrocytes. Consistent with our hypothesis, this cell cluster has an RNA velocity trajectory that moves inwards from both the erythrocyte and hepatocyte clusters (**Figure 1F**), indicating that these cells share markers with both cell types. Thus, we hereafter refer to this cluster as “efferocytosing hepatocytes”.

Together, we provide a cell atlas resource of key tissues across sexes in the turquoise killifish, with annotations for cell types found in these niches. The promoter sequences of top markers genes validated here in this species could ultimately be used to drive cell type specific transgene expression.

### Global analysis of sex differences across tissues and cell types

Since we profiled individuals of both sexes, we next analyzed our dataset to identify any potential large-scale sex differences: (i) at the level of transcriptional landscapes in each cell type, or (ii) in terms of the underlying cell composition of tissues (**Figure 2A**). First, there was no clear bias in median number of unique genes per cell type across cohort and sex (**Supplementary Figure S2A**). Next, we used AUGUR^44, 45^ to identify the cell type(s) transcriptionally most responsive to sex as a biological variable (**Figure 2B, C**). AUGUR identified hepatocytes among the most sex-responsive cell types. However, the most sex-dimorphic cell type was endothelial cells, which may be driven by direct exposure of endothelial cell to circulating sex hormones^46^. NK/T-cells and their progenitors also showed strong sex-dimorphism (**Figure 2B,C**).

**Figure 2.**
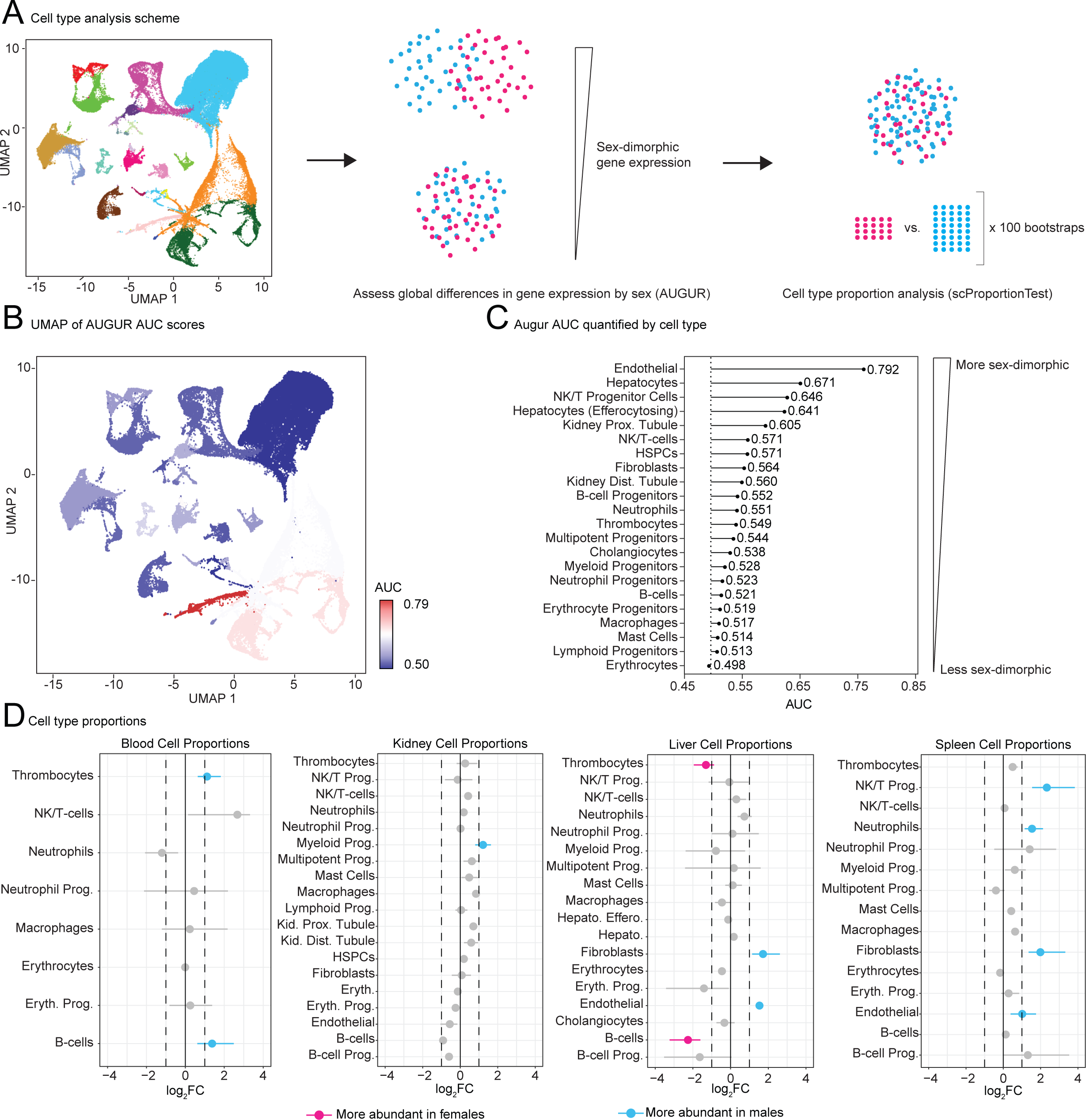
Sex-dimorphism in gene expression profiles and proportions across cell types in the African turquoise killifish. **(A)** Outline of cell type-level analysis pipeline. Cells were annotated according to cell types from the complete tissue atlas. The magnitude of gene expression differences between cell types of different sexes were analyzed by AUGUR and cell type proportions between sexes were analyzed using *scProportionTest*. **(B)** UMAP representation of AUGUR AUC results. Note that values close to 0.5 denote smaller sex differences (blue shades), and values closer to 1 reflect larger sex differences (red shades). **(C)** Lollipop plot of AUGUR AUC results per cell type. **(D)** Cell type proportion differences between sexes for each tissue. Left-shifted cell types are more abundant in females and right-shifted cell types are more abundant and males. The dashed lines represent significance thresholds of >1 log_2_ fold-change in cell abundance, <0.05 p-value.

We then examined the cellular composition of each tissue as a function of sex (**Supplementary Figure S2B**). As expected, blood was comprised almost entirely of erythrocytes, whereas other tissues showed more cell diversity. Unsurprisingly, hepatocytes and efferocytosing hepatocytes represented the bulk of liver cells, in both female and male tissues. The kidney contained a wide variety of blood cell types, consistent with its role as the primary hematopoietic organ in the turquoise killifish. The spleen also contained many immune cell types, but hosted a large proportion of erythrocytes, consistent with reports of the teleost spleen as a primary erythropoietic organ.

Next, to determine whether there were any significant differences in cell proportions for each tissue between sexes, we used the single-cell RNA-seq-specific scProportionTest tool^47^, which uses permutation testing to quantify cell type proportion changes between groups (**Figure 2D**). Compared to females, males showed a significant (>1 log_2_ fold-change, <0.05 p-value) enrichment for thrombocytes and B-cells in the blood; myeloid progenitors in the kidney; and NK/T-cell progenitors, neutrophils, fibroblasts, and endothelial cells in the spleen. In the liver, males were enriched for fibroblasts and endothelial cells while females showed an enrichment for B-cells and thrombocytes. Of note, the liver and blood showed mostly inverse cell type enrichment profiles between the sexes.

### Sex-dimorphic gene expression across cell types in the turquoise killifish

We next examined differential gene expression profiles in each cell type as a function of sex (**Figure 3A**). Since single-cell level differential expression suffers from high false discovery rates^48^, we used muscat^49^ to obtain pseudobulk gene expression profiles for each cell type and each cohort. This approach generates a conventional gene expression matrix, on which we could perform differential gene expression analysis for each cell type using DESeq2^50^ (**Figure 3A**). Importantly, to prevent technical noise from overshadowing biological effects, only cell types with at least 25 cells in both sexes for each of the 3 cohorts were further analyzed for differential expression analysis, yielding 13 cell types with robust pseudobulk expression profiles for downstream analysis.

**Figure 3.**
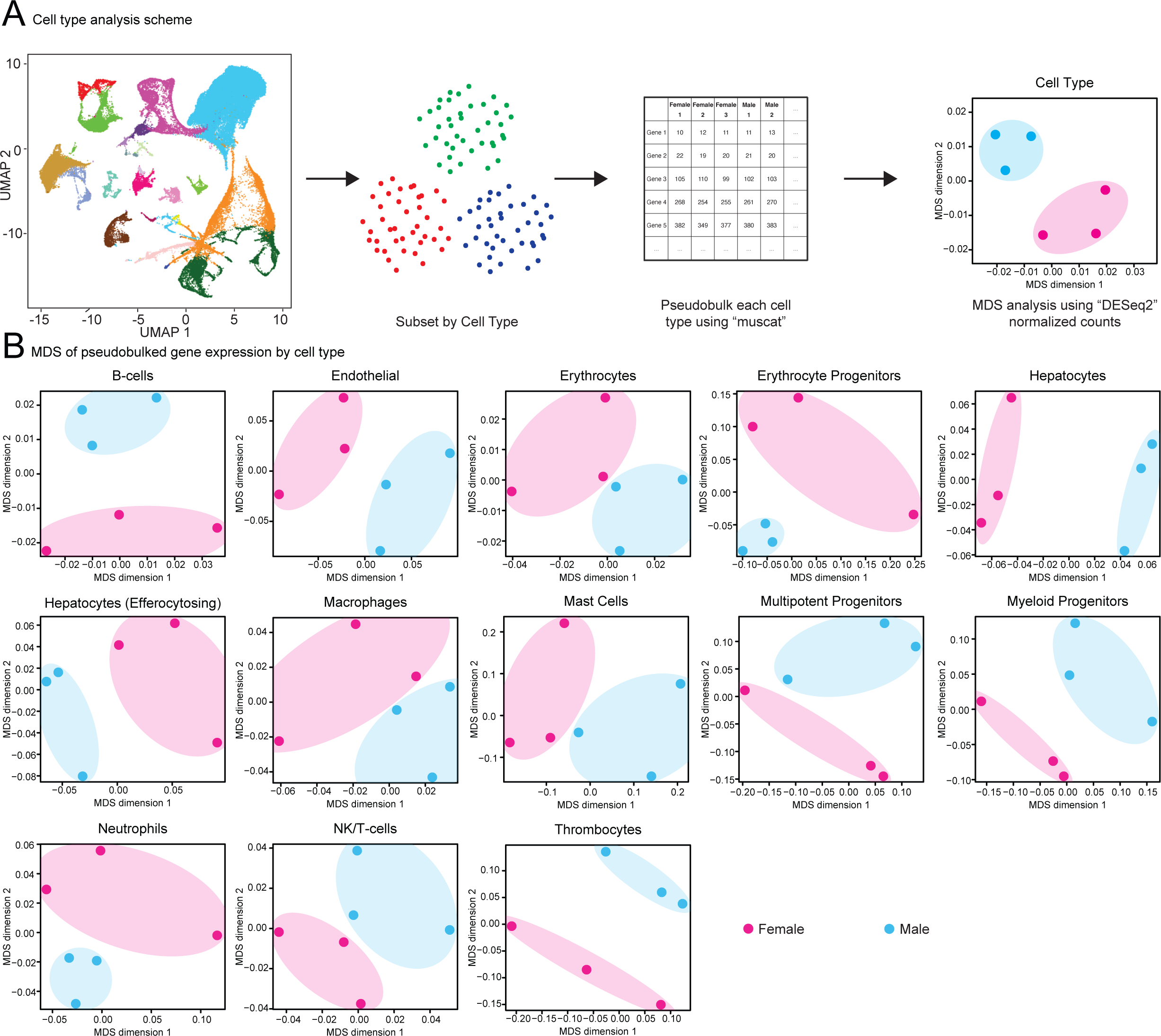
MDS analysis of pseudobulked cell type transcriptomes reveals widespread sex-dimorphism across cell types in the African turquoise killifish. **(A)** Schematic for differential gene expression analysis. Cell types were partitioned from the tissue atlas and their transcriptional profiles pseudobulked using muscat. Global gene expression differences for each cohort were assessed using MDS plots. **(B)** MDS plots for each cell type with at least 25 cells per sex in all 3 cohorts.

To visualize the overall similarity of transcriptomic landscapes across biological samples, we performed multi-dimensional scaling (MDS) analysis (**Figure 3A,B** and **Supplementary Figure S3A**). Importantly, replicates clustered more tightly by cell type than by sex (including hepatocytes), which is consistent with the fact that cell type has a larger effect on gene expression than sex. Strikingly, when performing MDS analysis for each cell type, male and female pseudobulk samples clearly separated, suggesting sex-dimorphic gene expression in all studied cell types (**Figure 3B**).

Next, we performed differential gene expression analysis as a function of sex across cell types to identify genes with significant sex-dimorphic gene expression (FDR < 5%; **Figure 4A**). Since both genes and TEs could show sex-dimorphic expression patterns, we analyzed genes and TEs together^26, 51^. We first compared global patterns of sex-dimorphism in gene expression by determining which cell types showed most correlated changes in gene expression as a function of sex using Spearman rank correlation Rho (**Supplementary Figure S4A**). We observed 3 major clusters with similar differential expression signatures: 1) multipotent progenitors, thrombocytes, B-cells; 2) neutrophils, erythrocyte progenitors, myeloid progenitors, mast cells; 3) hepatocytes, erythorocytes, NK/T-cells, efferocytosing hepatocytes. While most cell types had correlated gene expression profiles, macrophage and B-cell gene expression profiles were negatively correlated with those of hepatocytes, erythorocytes, NK/T-cells and efferocytosing hepatocytes.

**Figure 4.**
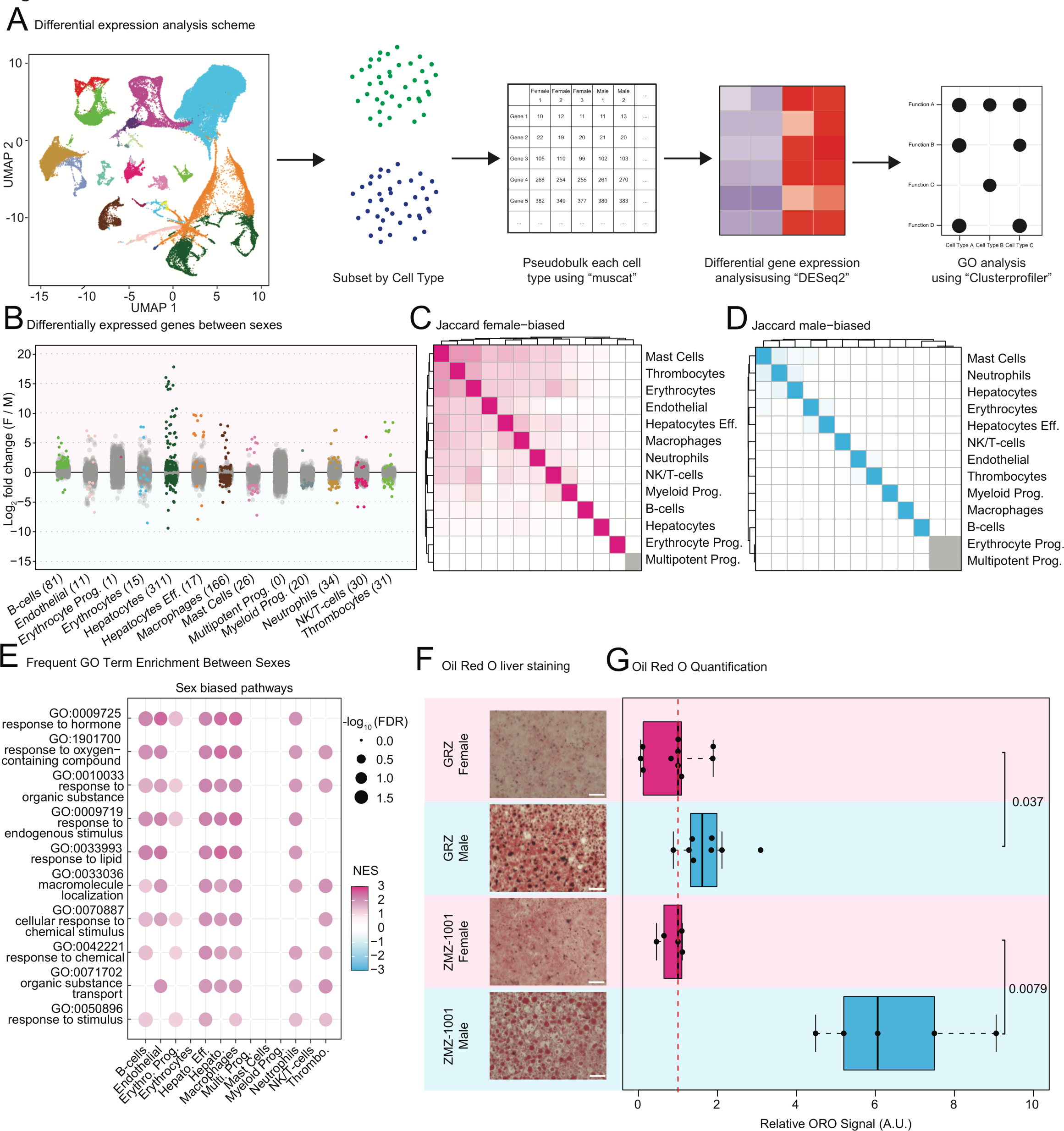
Differential gene expression analysis reveals sex-dimorphism in lipid metabolism pathways across cell types in the African turquoise killifish. **(A)** Overview of differential gene expression analysis and Gene Ontology (GO) analysis by cell type. Cell types was partitioned from the completed tissue atlas and analyzed for differential regulation using pseudobulked muscat analysis. Differential gene expression analysis was performed with DESeq2 and GO analysis was performed with ClusterProfiler. **(B)** Strip plot of differentially expressed genes per cell type (FDR < 5%). Differentially expressed transcripts are highlighted while non-significant genes are in gray. Transcripts above the midline are upregulated in females while transcripts below the midline are enriched in males. **(C-D)** Jaccard plots of female-biased (**C)** and male-biased **(D)** transcripts by cell type. Female cell types tend to share a more common set of upregulated genes than males. **(E)** GO terms enriched in at least 6 cell types. Note that all shared terms were enriched in females. **(F)** Representative images of Oil Red O staining of GRZ and ZMZ1001 livers from both sexes. Note that male livers, regardless of strain, tend have a larger stained area compared to females. Scale bar = 50 µm. **(G)** Quantification of Oil Red O staining in 6-week-old turquoise killifish livers. Percentage of stained area was obtained for each sample. To mitigate batch-to-batch staining intensity variation, all values are normalized to the median value of the females of the cognate strain in a batch. Significance in Wilcoxon rank sum test are reported.

We then examined which transcripts were significantly differentially expressed between sexes (FDR < 5%; **Supplementary Table S4**). At the cell type level, hepatocytes unsurprisingly had the highest number of sex-dimorphic transcripts (311) of any cell type, followed by macrophages with 166 transcripts (**Figure 4B**). Interestingly, female-biased transcripts were often upregulated to a larger magnitude than male-biased counterparts (**Figure 4B; Supplementary Table S4**). Further, although most sex-dimorphic transcripts were unique to each cell type, female-biased transcript were shared to a greater extent than male-biased transcripts across cell types, as measured by Jaccard index (**Figure 4C,D**). This observation is consistent with the existence of a core common female transcriptional signature across turquoise killifish cell types.

We next asked which genes tended to be sex-dimorphic most frequently across cell types (**Supplementary Figure S4B**; **Supplementary Table S4**). The most frequently female-biased genes were *LOC107388898* (hereafter referred to as *ncFem1*), *zp3* and *vitellogenin-1-like* (in 10, 9, and 9 cell types, respectively; **Supplementary Figure S4B**). The most frequently male-biased genes were *hpx*, *apoa1a*, and *aldob* (in 5, 4, and 3 cell types, respectively; **Supplementary Figure S4B**). We independently confirmed the upregulation of the most highly male-biased transcripts, *hpx*, by RT-qPCR in liver, spleen, and muscle tissue (**Supplementary Figure S4C**). We observed significant male-bias in *hpx* expression in liver and spleen tissues (p ≤ 0.01 in non-parametric Wilcoxon test), and a non-significant trend for upregulation in muscle (p ∼ 0.07 in non-parametric Wilcoxon test), likely due to large inter-individual variability (**Supplementary Figure S4C**). We also validated female bias of our top female-biased transcript *ncFem1* in the same sample set (see details in next section; **Figure 5F**). This suggests that sex-specific gene expression patterns identified in our atlas could be predictive of sex-biased gene expression in tissues even outside of the scope of our atlas.

**Figure 5.**
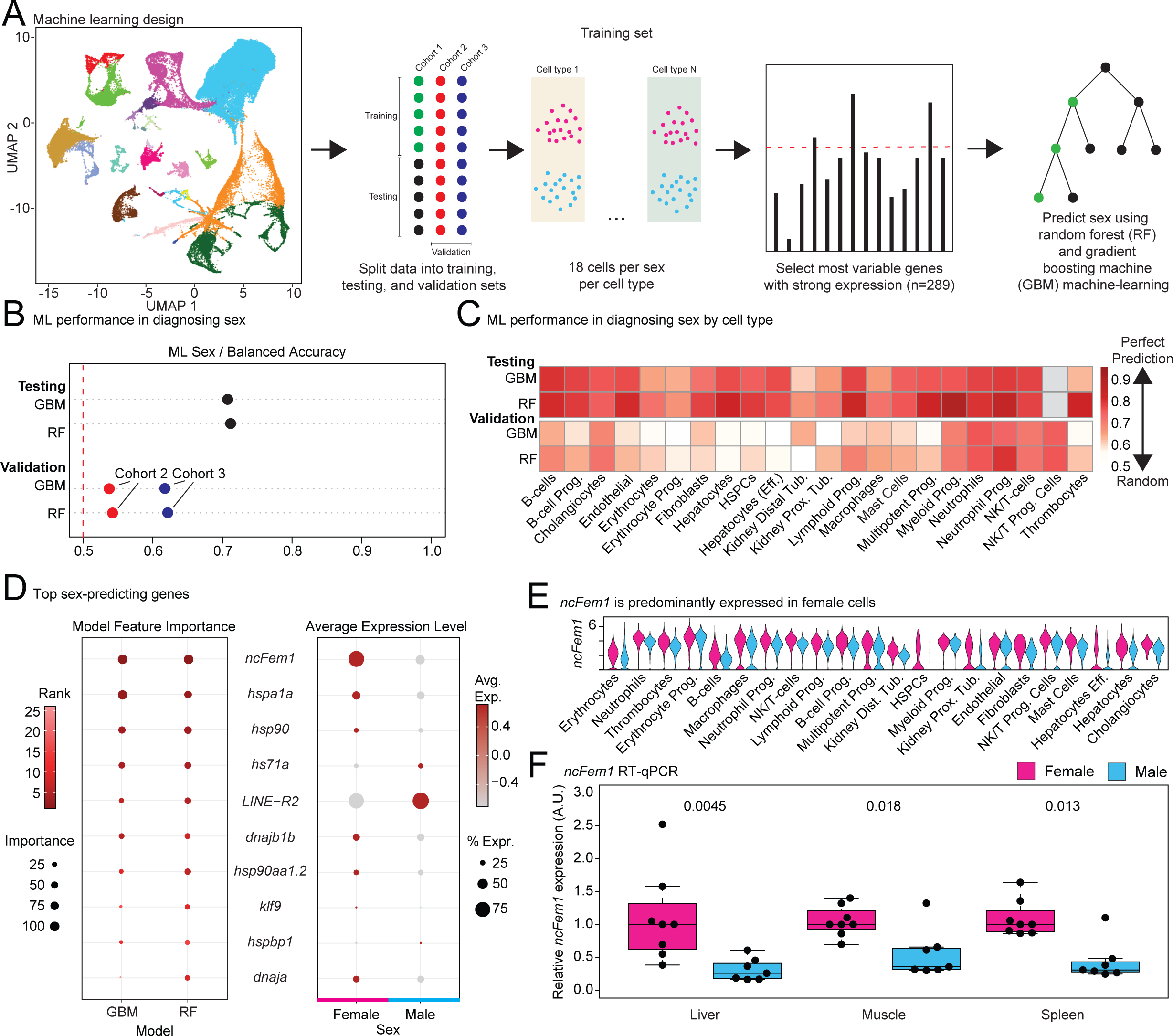
Machine-learning identifies novel sex-specific transcriptional markers in African turquoise killifish cells and tissues. **(A)** Machine learning schematic. Data was split into training (cohort 1), testing (cohort 1), and validation (cohorts 2 and 3) sets. For the training set, 18 cells per cell type per sex were used. All other cells from cohort 1 correspond to the testing set. The most variable genes from the training set that were robustly expressed across cell types were used to train Random Forest (RF) and Gradient Boosted Machines (GBM) machine-learning models to predict sex. **(B)** Performance of machine-learning models on the testing and validation datasets. Note that RF tends to outperform GBM slightly. **(C)** Machine-learning performance in determining sex segregated by cell type on the testing and validation sets. RF tends to outperform GBM across cell types. **(D)** The top 10 sex-predicting transcripts from both machine-learning models (left) and their average expression across the entire tissue atlas by sex. The most sex-predictive transcripts show strong sex-dimorphism. Since many NCBI gene names are uninformative, names corresponding to their homologs are used. Original NCBI gene names are provided in **Supplementary Table S9**. **(E)** Violin plots of the top female predictive transcript (*ncFem1*) in each cell type. In each cell type, *ncFem1* is more highly expressed in females. **(F)** Validation of the female-biased expression for *ncFem1* by RT-qPCR in liver, muscle, and spleen tissue. In each tissue, *ncFem1* is more highly expressed in females. Significance in Wilcoxon rank sum test are reported.

To further understand which functional processes might be impacted by sex-dimorphic gene expression, we performed a gene set enrichment analysis (GSEA) on each cell type querying for Gene Ontology (GO) terms in the “biological process” category (**Figure 4A**) and reported the most significantly enriched GO terms for each cell type, if any (**Supplementary Figure S4D; Supplementary Table S5**). To determine which biological processes were commonly differentially regulated between sexes, we also identified the top 10 pathways that were significantly sex-dimorphic according to GSEA in ≥ 6 different cell types (**Figure 4E**). Intriguingly, common pathways were exclusively enriched in female cells. One of the most enriched terms in females was “response to hormone”, suggesting that the female hormonal milieu may drive gene expression differences across somatic cell types. Such a gene expression response is consistent with the higher degree of shared female-biased genes as measured with our Jaccard index analysis (**Figure 4C**).

Another GO term commonly enriched in female cells was “response to lipid” (**Figure 4E, S4D**). Fish, including the African turquoise killifish, generally accumulate energy storage molecules such as glycogens and lipids in the liver^52–55^. Therefore, we analyzed turquoise killifish livers for sex-dimorphism in lipid storage using Oil Red O staining on fresh liver tissues of young adult killifish. Interestingly, we observed clear sex-dimorphism in the lipid content of male and female livers from the short-lived inbred GRZ turquoise killifish strain, which was used in this study for transcriptomic profiling (**Figure 4F,G**). To exclude the possibility that the high lipid content of male livers was due to a mutation in the GRZ inbred strain, we also assessed lipid deposition patterns in the livers of young adult turquoise killifish from the longer-lived ZMZ-1001 wild-derived strain, which revealed an even starker sex-differences in lipid storage patterns (**Figure 4F,G**). Thus, we believe that turquoise killifish, regardless of strain, experience stark sex-dimorphism in lipid storage and utilization, consistent with observed sex-dimorphic transcriptional signatures.

### Identifying potential molecular markers of sex using machine-learning

Since we observed sex-dimorphic gene expression, we next asked whether the sex of a turquoise killifish cell could be accurately predicted from gene expression using machine-learning approaches (**Figure 5A**). For this purpose, we split cells from our atlas into training, testing, and validation sets (see Methods). Training and testing were performed on cohort 1. To limit issues due to unequal representation of cell types in the training set, we used a relatively small training set, randomly sampling 18 cells per cell type per sex (since the cell type with the smallest representation per sex in this cohort, NK/T progenitor cells, was only comprised of 18 female cells). All cells from cohort 1 not used for training were used for model testing. From the training data, we identified 289 genes that were highly variable but robustly expressed across at least 20 cell types (*i.e*. genic features), which were used to train random forest (RF) and gradient boosting machine (GBM) models (see Methods; **Figure 5A**). Accuracy of models were tested on withheld cells from cohort 1, and then validated using cohorts 2 and 3.

Both model types performed similarly well at predicting sex (**Figure 5B; Supplementary Figure S5A**). The testing balanced accuracy for each model was ∼70% (**Figure 5B**). GBM and RF models were approximately >50% and >60% accurate in predicting sex in cohorts 2 and 3 across all cell types, respectively, with a slight edge for the RF model (**Figure 5B**). When segregated by cell type, predictive accuracy varied widely with generally poor performance for erythrocytes and much better performance for immune cell types (**Figure 5C**). The poor performance of the erythrocytes is consistent with having the lowest sex-dimorphic score with AUGUR (**Figure 2B,C**).

We next took advantage of the models to ask which features had the most weight in driving model accuracy (**Figure 5D** left panel; **Supplementary Table S6**). As expected, the top genic features tended to show sex-biased gene expression intensities (**Figure 5D** right panel, **Supplementary Figure S5B**). In both models, the top predictor for females was a non-coding RNA, *LOC107373896*, which we have renamed *ncFem1* due to systematic female-bias across cell types (**Figure 5D; Supplementary Figure S4B** and **S5B**). The top male predictive transcript was *hs71a*, though it was not expressed in all cell types (**Figure 5D, Supplementary Figure S5B**). The most sex-predictive male transcript detected in all cell types was a TE, *LINE-R2* (**Figure 5D; Supplementary Figure S5B**).

To confirm whether high expression of *ncFem1* and/or *LINE-R2* can generally be used as markers of sex, we performed RT-qPCR for *ncFem1* (**Figure 5F**) and *LINE-R2* (**Supplementary Figure S5C**) on two tissues used in our atlas, spleen and liver. We also sampled muscle (a tissue not included in our atlas) to determine if our approach could detect sex-biased genes more broadly (**Figure 5F; Supplementary Figure S5C**). We identified a strong and significant female-bias for *ncFem1* expression across all assayed tissues (**Figure 5F**). However, we did not find any such enrichment for *LINE-R2* in male samples (**Supplementary Figure S5C**). Although we are unsure as to why sex-bias could not be confirmed, we surmise this may result from the multi-copy nature of the element, sequence variation between copies, and potential differential impact on expression from intronic *vs*. autonomous copies^56^. However, lack of robust sex-dimorphism for *LINE-R2* by RT-qPCR is consistent with it not being identified as a top recurrent sex-biased gene with our DESeq2 analysis (**Supplementary Figure S4B; Supplementary Table S4**).

### Using the turquoise killifish cell atlas to perform bulk tissue RNA-seq deconvolution

Single-cell datasets can be used to deconvolute bulk RNA-seq datasets and determine underlying cell composition^26, 57–59^. Thus, we asked whether our atlas could be used to uncover new biology from published bulk RNA-seq datasets.

First, we attempted to use our killifish atlas to deconvolute a recently published killifish bulk RNA-seq dataset from McKay *et al.* ^60^. McKay *et al*. generated liver transcriptomes from male and female GRZ 9-week-old turquoise killifish fed *ad libitum* or calorically restricted diets. We deconvoluted the dataset using liver cells from our atlas using the deconvolution program CSCDRNA^61^ (**Figure 6A**). We first compared the overall average predicted cell type composition of each biological group from McKay *et al.* (*ad libitum* fed female livers, calorically restricted male livers, etc.) to male and female liver cell type composition from our cell atlas (**Supplementary Figure S6A**). Overall cell type composition is similar with large fractions of hepatocytes, efferocytosing hepatocytes, and erythrocytes. Next, we quantified the effects of sex and caloric restriction on liver cell type composition. When comparing deconvoluted male and female livers from both diet groups, we found a significant enrichment in B-cells and macrophages in females and in erythrocyte progenitors in males (**Figure 6B; Supplementary Figure S6B**). This agrees with our analysis of cell type proportions in **Figure 2D** which shows an enrichment of B-cells in the female liver. Next, we assessed whether caloric restriction had an impact on cell proportions in the liver, in males and females separately (**Supplementary Figure S6C**). At least in this dataset, we could not find significant evidence for large scale changes in underlying cell type composition in response to this intervention.

**Figure 6.**
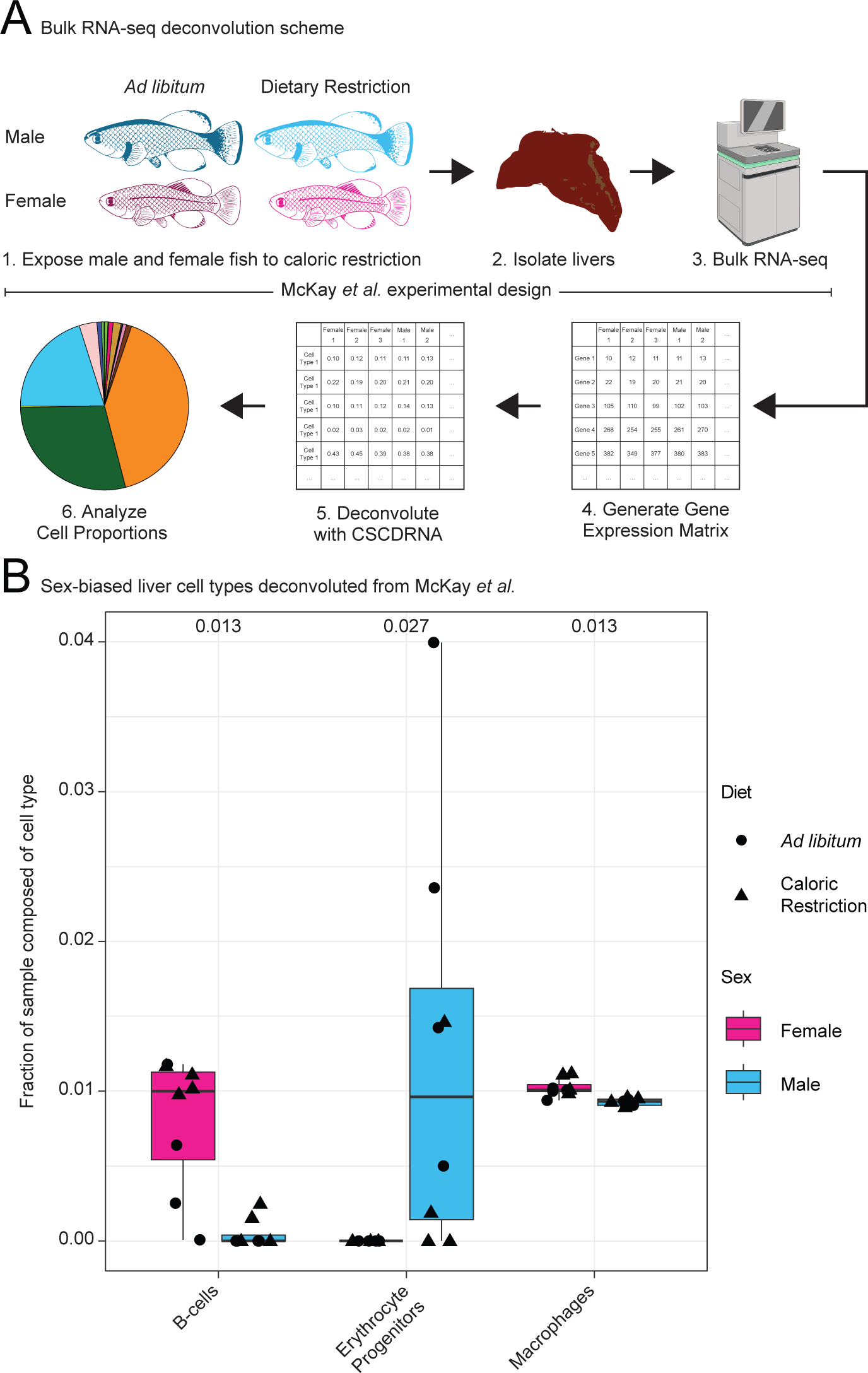
The African turquoise killifish annotated single-cell atlas can be used to infer cell type proportions in bulk RNA-seq datasets. **(B)** Outline of the deconvolution process. Publicly available bulk RNA-seq files from McKay *et al.* were downloaded, mapped to the killifish genome, and counted over genes. Count matrices were deconvoluted using CSCDRNA using liver data from the tissue atlas to generate cell type proportion estimations from each RNA-seq library. **(B)** Boxplot of B-cells, erythrocyte progenitors, and macrophages from male and female livers from fish from McKay *et al*. For each cell type, the cell type proportions from each sex were contrasted using a Wilcoxon-Rank sum test with Benjamini-Hochberg p-adjustment for multiple testing. Both dietary groups were included, and are coded by point shape. Shown cell types had significantly different predicted abundances between males and females. All cell types shown in **Supplementary Figure S6B**. Significance values for all cell types can be found in **Supplementary Table S8**. B-cells and macrophages are significantly enriched in females and erythrocyte progenitors are significantly enriched in males. Females: pink, Males: blue, *ad libitum* diet: circle, calorically-restricted diet: triangles.

## Discussion

In this study, we generated and characterized the first multi-tissue single-cell atlas for the African turquoise killifish profiling blood, kidney, liver, and spleen. Our atlas includes data from both sexes, which is crucial for detecting sex-specific and conserved aspects of turquoise killifish biology. We were able to define 22 distinct cell types and clean marker genes for these cell types, which should have great utility to the growing turquoise killifish research community.

Furthermore, through this analysis, we were able to uncover interesting features of killifish biology. First, we observed that, as expected for a teleost fish, most hematopoiesis occurs in the kidney and erythropoiesis in the spleen. Furthermore, we identified a new abundant cell state in the killifish liver in the form of efferocytosing hepatocytes. Although efferocytosis by hepatocytes had been previously observed^43^, our study is the first report of such a phenomenon in this species. This finding suggests that a major role of turquoise killifish hepatocytes may be to clear dead and/or dying erythrocytes from circulation.

Overall, we observed the strongest evidence of sex-dimorphic biology in the turquoise killifish liver. Sex-dimorphism in liver transcriptomes has also been observed in other species, including mouse ^62^ and humans^63^. Not only was sex-dimorphism observed at the transcriptional level, but we also observed stark differences in lipid storage in liver tissue. This could indicate a hormonally-driven program of differential lipid utilization. It is tempting to speculate that vitellogenin, which is produced by females and is part of the large lipid transfer protein (LLTP) superfamily, could facilitate the differential lipid utilization and export from the liver^64^. Importantly, in contrast to female livers, male livers contained many lipid droplets. Although fatty livers are known to be common even in wild killifish^55^, the stark contrast in lipid storage in the livers of female vs. male killifish may provide an interesting handle to study fatty liver disease, a disease commonly associated with aging^65^, and is therefore, a promising area for future research.

Our atlas will be a useful resource for the killifish research community. We demonstrated that machine learning can be used to successfully predict sex and identify sex-biased genes. These genes could be used to validate the sex of samples, for instance as quality control checks in bioinformatic pipelines. In addition, although we focused on sex-differences in this study, our data could also be used to identify new unexplored features in the killifish immune and metabolic systems. Finally, using a proof-of-principle analysis, we showed that our atlas can be used for bulk RNA-seq deconvolution by demonstrating that cell proportions can be reliably estimated from publicly available tissue-level bulk RNA-seq data.

In summary, we present a multi-tissue, multi-sex atlas of aging-relevant tissues in the killifish that should drive hypothesis testing for an emerging aging model and enable new sophisticated analyses in this model.

## Materials and methods

### African turquoise killifish husbandry

African turquoise killifish were raised according to gold standard procedures ^66^. To decrease risks of aggression, adult fish were single housed in 2.8L tanks on a recirculating aquatics system manufactured by Aquaneering Inc. System water parameters were as follows: temperature: 29°C; pH: 7.3; conductance: 670-750 μS; ammonia and nitrite: 0ppm and Nitrate: ∼15 ppm. Adult fish were fed twice per day with Hikari Freeze Dried Bloodworms 4-6 hours apart, and live *Artemia* once per day. Fry were reared in system water incubated at 28°C, then placed on the recirculating system starting at 2 weeks post hatch. Fry were fed live *Artemia* exclusively until 4 weeks post hatch. The fish facility was kept on a light/dark cycle of 13/11 hours (lights on 9am-10pm). The fish were euthanized by immersion in 1 g/L of Tricaine MS-222 dissolved in system water followed by decapitation. All fish were euthanized between 10 am and 12 pm to minimize circadian effects. All husbandry conditions and experimental procedures were approved by the University of Southern California IACUC. Animal care and animal experimentation were performed in accordance with IACUC approved protocols for *Nothobranchius furzeri at* the University of Southern California (approved protocols 21223 and 21215).

### Tissue dissection and dissociation

Three cohorts of 6-week-old GRZ strain *N. furzeri* were used to create three sets of 10xGenomics single-cell libraries. For the first cohort, cells were pooled from 4 female and 3 male fish to ensure that enough cells would be isolated from the generally smaller female tissues. Since more than sufficient cell totals were obtained from the first female pool, the next two cohorts consisted of 3 female and 3 male fish. For each cohort, tissues from different fish of the same sex were pooled and dissociated together to maximize viability and cell yield. In each cohort, blood, liver, kidney, and spleen were obtained. All kidney tissue (head and tail kidney) was taken. Except for blood, all dissected organs were kept on ice in Dulbecco’s Modified Eagle Medium (DMEM) (Corning, 15-013-CV) until all dissections were complete, which took up to 1 hour.

Blood was obtained by cutting off the tail of freshly euthanized fish roughly 1-2 mm into the vascularized caudal peduncle tissue. A drop of 1 mM EDTA was then deposited on to the cut site to prevent blood coagulation. Fish were then held with the tail oriented downwards to produce a drop of blood by gravity flow. If a drop of blood was not produced by gravity, fish were gently squeezed. If blood was not produced on the original attempt, a second cut was made anterior to the previous cut site and the process was repeated. A single drop of blood (<= 10 µL) was collected using a low retention pipette tip (USA Scientific, 1183-1710) pre-coated in 1 mM EDTA and deposited into a 5 mL ice-cold of PEB buffer (PBS, pH 7.2, 0.5% BSA, 2 mM EDTA; dilution of MACS BSA Stock Solution [Miltenyi Biotec, 130-091-376] 1:20 with autoMACS® Rinsing Solution [Miltenyi Biotec, 130-091-222]).

All other solid organs were dissociated into single-cell suspensions using Miltenyi’s Liver Dissociation Kit (Miltenyi Biotec, 130-105-807) with the gentleMACS Octo Dissociator with Heaters (Miltenyi Biotec, 130-096-427) according to manufacturer’s instructions. Cell viability was assessed on a Countess II FL Automated Cell Counter (Thermo Fisher Scientific, AMQAX1000) using a 1:1 dilution of Trypan Blue Stain (Thermo Fisher Scientific, T10282) on Countess Cell Counting Chamber Slides (Thermo Fisher Scientific, C10228). Due to initially low cell viability (<50%), liver samples in cohort 2 were reduced of dead cell burden using the EasySep Dead Cell Removal (Annexin V) Kit (STEMCELL Technologies, 17899) to improve cell viability to >70%. For the third cohort, all samples were preemptively subjected to treatment with the Dead Cell Removal Kit. Samples excluded for low viability were the female spleen sample from cohort 2 and both blood samples from cohort 3 (**Supplementary Figure S1B**). Although sufficient cells were obtained, a wetting failure led to unsuccessful library construction for cohort 2 male kidney (**Supplementary Figure S1B**).

Singlet rate was estimated on dissociated tissues using flow cytometry on a MACSQuant Analyzer 10 (Miltenyi Biotec, 130-096-343). Flow cytometry data was analyzed using Flowlogic v8 (Miltenyi Biotec, 160-002-087). Cells were identified using a common gate on FSC-A *vs*. SSC-A, and singlets were assessed by evaluating the linear grouping of cells by FSC-A *vs*. FSC-H. Our dissociation protocol yields good dissociation rates across the three solid tissues as measured on one set of samples (>95% singlets, **Supplementary Table S1**).

### Single-cell library preparation

Single-cell libraries were prepared using Chromium Next GEM Single Cell 3L GEM, Library & Gel Bead Kit v3.1 (10X Genomics, PN-1000121) according to manufacturer’s instructions (10xGenomics User Guide Chromium Next GEM Single-cell 3′ Reagent Kits v3.1 (CG000204, Rev D)). Based on Countess estimates, cell suspensions were loaded for a targeted cell recovery of 7,000 cells per sample. Samples were run on Chromium Next GEM Chip G (10X Genomics, 2000177) per manufacturer’s instructions. Partially amplified and final single-cell RNA-seq libraries were quantified by a dsDNA Quantitation, high sensitivity assay (Invitrogen, Q32854) on a Qubit® 3.0 Fluorometer (Thermo Fisher Scientific, Q33216). Completed single-cell libraries were assessed for quality on the 4200 TapeStation system (Agilent Technologies; G2991A) with a High Sensitivity D1000 DNA ScreenTape (Agilent Technologies 50675584). Libraries were sequenced on an Illumina Novaseq 6000 generating 150 bp paired-end reads at Novogene USA. Raw FASTQ reads have been deposited to the sequence read archive under accession PRJNA952805.

### BLAST identification of mouse and zebrafish homologs for annotation

We obtained predicted turquoise killifish protein sequences for the version of the genome GCF_001465895.1 from NCBI. Mouse protein sequences were obtained from Ensembl Biomart version 107, which contains annotations derived from genome version GRCm39 (accessed 2022-10-10). Zebrafish protein sequences were obtained from Ensembl Biomart version 105, which contains annotations derived from genome version GRCz11 (accessed 2022-02-11).

In both cases, NCBI BLAST 2.10.0+ was used to align the other species proteomes to the killifish reference proteome to obtain the best hit with -best_hit_score_edge 0.05 and - best_hit_overhang 0.25. To reflect the relative phylogenetic distance, an e-value cutoff of 1e-3 was used for mouse homology, and 1e-5 for zebrafish homology.

### 10X Genomics data processing

The following pipeline was run for each set/cohort of single-cell datasets. A custom genome reference was assembled using the GCF_001465895.1 genome assembly ^67^, which has been annotated by NCBI. The genome was hardmasked using a turquoise killifish repeat library from FishTEDB ^68^ (accessed 2021-09-06). TE sequences were then added as independent contigs to the genome reference fasta and as entries to the gene reference file gtf. CellRanger 6.1.2 (10xGenomics) was used to create the reference.

FASTQ read files were aligned to this turquoise killifish reference genome and quantified using “cellranger count” (CellRanger 6.1.2). Ambient RNA was removed using the SoupX ^69^ package in R v3.6.3^70^ using the “out” folder from cellranger. Quality control was performed in R v3.6.3 using Seurat v3.2.2^71–74^ by removing dead and low-quality cells by selecting for cells that contained between 250-7000 UMIs and <30% mitochondrial reads.

A list of mouse cell cycle genes was obtained from the Seurat Vignettes (https://www.dropbox.com/s/3dby3bjsaf5arrw/cell_cycle_vignette_files.zip?dl=1), derived from a mouse study^75^. Cell cycle phase was predicted using homologous turquoise killifish cell cycle genes, and using the function CellCycleSorting() to assign cell cycle scores to each cell. Multiplets were annotated using a combination of Doubletfinder v2.0.3^76^ and scds v1.2.0^77^ for each particular 10xGenomics library. Only cells called as singlets by both methods were used to create the cell atlas.

After pre-processing, the 3 cohort datasets were merged into a single Seurat object. Gene expression was then normalized on a global scale using SCTransform v0.3.2^78^, regressing to the variables “nFeature_RNA”, “percent.mito”, and “Batch” with the parameter variable.features.n = 5000. Reciprocal PCA was used to integrate data from the 3 cohorts and mitigate batch effects, using the top 2000 most variable genes and k = 20.

### Cell type annotation and AUGUR

We used Seurat to perform unsupervised cell clustering with a single nearest neighbor resolutions and a resolution of 0.8, which was chosen to provide granularity pre-annotation with 50 clusters. This clustering served as the basis for following annotation efforts.

Four different R packages, scSorter v0.0.2^79^, code adapted from scType v1.0^80^, CellAssign v0.99.21^81^ (marker-based) and SingleCellNet v0.1.0^82^ (cross-species atlas-based annotation) were used to obtain first pass predictions of cell type identity. For marker-based annotation methods, published cell type markers from a wide array of zebrafish cell atlases were used^29–37^ to obtain a list of candidate markers (see above**; Supplementary Table S3**), that were then converted to their turquoise killifish homolog in R, if any, using turquoise killifish homologs of zebrafish genes (see above). SingleCellNet v0.1.0 was trained using data from the Tabula Muris reference dataset^83^ using turquoise killifish homologs of mouse genes (see above). Cells were labeled as a particular cell type when 3 out of 4 automated approaches agreed on cell type. Any remaining unannotated clusters were annotated manually by taking into consideration marker genes according to Seurat compared to other clusters, tissue of origin, cell cycle score, and nearest neighbors in gene expression space. When two clusters shared expression of typical markers, clusters with high proportions of proliferative cells were annotated to more progenitor states (**Figure 1D**).

Once all cell types were annotated, we used the R package AUGUR v1.0.3^44^ to determine which cell types were most transcriptionally sex-dimorphic.

### RNA velocity

RNA velocity^38^ was performed in loose accordance with the guide provided at https://github.com/basilkhuder/Seurat-to-RNA-Velocity. In a custom python v3.8.12 environment, the “out” folder of CellRanger was used to generate loom files containing single-cell data using velocyto v0.17.17^38^. Meanwhile, metadata describing cell identifiers, UMAP coordinates, and cell type was extracted from the completed atlas Seurat object. Loom files were converted into anndata files, appended with the appropriate metadata, then merged into a single anndata object. RNA velocity was performed on the merged anndata object using scvelo v0.2.5 ^84^.

### Cell Type Proportions Between Sexes

Cell type proportions were calculated from the final Seurat object using the R program scProportionTest v0.0.0.9000^47^ in R v4.2.2. Briefly, this program runs a permutation test on two different single-cell groups for each cell type and returns the relative change in cell type abundance between groups with a confidence interval for each comparison. Sexes were compared at the level of the entire tissue atlas as well as at the tissue level.

### Pseudobulk analysis for differential gene expression

Pseudobulking and differential gene expression were performed using R v3.6.3. The package muscat v1.0.1^49^ was used to pseudobulk the single-cell gene expression data by sex and cell type for downstream analysis. Only cell types with at least 25 cells across each sex/cohort were used to minimize technical noise for downstream analyses. The package RUVSeq v1.20.0 ^85^ was used on the pseudobulked object to remove noise linked to batch and tissue effects on gene expression. DESeq2 v1.26.0^50^ was used for differential gene expression analysis using the cleaned gene counts matrices from RUVSeq. Differential gene expression analysis was performed for genes and transposable elements (TEs). Differential gene expression results were used to determine the number of differentially expressed genes in each cell type by sex (**Supplementary Table 4**). We compared genome-wide log_2_ fold change as a function of sex across cell types using Spearman Rank correlation, and Jaccard indices were calculated to determine the degree of sharing for sex-biased genes across cell types.

### GO analysis for differential gene expression

The results from the differential gene expression were used as input for ClusterProfiler v3.14.3^86^ to run Gene Ontology enrichment using gene set enrichment analysis (GSEA). Zebrafish Gene Ontology terms were obtained from annotation package org.Dr.eg.db_3.10.0^87^. The DESeq2-derived t-statistic was used to rank genes for sex-biased gene expression, and turquoise killifish homology to zebrafish genes was used for the analysis (see above). Only “biological process” (BP) terms were considered for this analysis. We used 1000 permutations, minimum gene set size of 25 and maximum gene set size of 5000 to compute enrichment. Terms with a False Discover Rate < 0.05 were considered significantly sex-dimorphic (**Supplementary Table 5**).

### Machine-learning of sex markers

Machine learning was used to identify sex-predicting genes across cell and tissue types using R v3.6.3. To prepare data for machine learning, cohort 1 was set aside for training and testing data while cohorts 2 and 3 were used as validation sets.

Training and testing were performed on cells from cohort 1. To limit issues due to imbalanced representation of cell types, we used a relatively small training set, randomly sampling 18 cells per cell type per sex (since the cell type with the smallest representation per sex in this cohort, NK/T progenitor cells, was only comprised of 18 female cells). All cells from cohort 1 not used for training for were for model testing. From the training data, we identified 289 genes that (i) were among the top 5000 most variable genes in this subsetted dataset, and (ii) were expressed in at least 20 out of the 22 detected cell types, to be used as features in our machine-learning models. In addition, we also collected 4 other potentially important covariates (*i.e*. sex, tissue of origin, annotated cell type, and percent mitochondrial reads). The 289 genes and 4 covariates were used as training features.

To perform machine learning, we used the R package caret v6.0-86 to run the training step. Random forest (RF) and gradient boosting machine (GBM) were based on caret’s usage of packages randomForest 4.6-14 and gbm 2.1.8, respectively. For both model types, training was performed using 10-fold cross-validation to search a parameter grid. The models were selected to maximize balanced accuracy across female and male cells. Out-of-bag predictions across the 10 training folds of the best parameters are reported in **Supplementary Figure S5A**.

To determine testing and validation accuracy, withheld data from cohort 1 and data from Cohorts 2 and 3 were used, respectively. The caret ‘predict’ function was used to predict cell sex using the trained models. Prediction accuracy was determined globally by sex (**Figure 5B**), and for each individual cell type (**Figure 5C**). Variable importance scores were extracted by caret for the RF and GBM models. A ‘consensus’ ranking of top contributing variables was computed using the rank products of variable importance from each individual model (**Supplementary Table S6**).

### RT-qPCR validation of sex-dimorphic genes

Frozen liver, muscle, and spleen tissue from 6-week-old female and male GRZ fish were used for RNA extraction, cDNA synthesis, and quantitative PCR (qPCR). Each individual tissue was transferred to a Lysing Matrix D tube (MP, 6913500) with 1 mL of TRIzol (Ambion, 15596018) and homogenized using a Beadbug 6 microtube homogenizer (Benchmark Scientific, D1036) at 3,500 rpm for 30 seconds for 3 cycles. RNA was isolated according to the manufacturer instructions (Ambion, 15596018), with the addition of 1 µL Glycoblue reagent (Thermo Fisher Scientific, AM9516) as a carrier to improve RNA pelleting and recovery.

The cDNA was synthesized using the Thermo Scientific maximaTM H Minus cDNA Synthesis Master Mix (Thermo Scientific, MAN0016392) in the Thermal cycler C1000 (Bio-Rad, 1851196). qPCR was performed using the SensiFAST SYBR® No-ROX Kit (Bioline, BIO-98020) in the Magnetic Induction Cycler (MIC) machine (Bio Molecular Systems, MIC-2). The Cq values were obtained using the micPCR v2.12.7 software, with a dynamic threshold setting. To normalize expression values, we used amplicons for 3 housekeeping genes (the turquoise killifish homologs of *hprt*, *actb*, *sdha*) to minimize technical noise and used the geometric mean of normalizing amplicons as the value of reference for the sample. Relative gene expression was obtained with the delta-delta Ct method, normalized to the median value of females in a tissue of interest for each set of samples. Significance was tested by Wilcoxon test. Primers were designed using Primer3 software, selecting for amplicon lengths of 100-200 bps^88^. Primers are listed in **Supplementary Table S7**.

### Liver histology and lipid content analysis

Livers from euthanized male and female 6-week-old GRZ and ZMZ1001 turquoise killifish that had been fasted for 18 hours prior to euthanasia were harvested and fresh frozen on dry ice. Frozen livers were mounted in Tissue-Tek O.C.T Compound (Sakura, 4583) and sliced into 13 µm slices using PTFE-coated microtome blades (Duraedge, 7223) at -10^°^C on a Cryostat CM1860 (Leica, 14-0491-46884). Liver sections were mounted on VWR Superfrost Plus slides (VWR, 48311-703) and stained with 0.5% Oil Red O (Sigma 00625-100G) in 60% Triethyl Phosphate (Sigma 538728-1L). The slides were preserved with Glycerol Gelatin (Sigma GG1-15ML). Liver slices were imaged on a Keyence BZ-X710 at 40x magnification.

Slices were quantified using ImageJ v1.54d. Per animal, 2 images from 2 independent liver slices were processed (4 images per animal). Briefly, the images were converted from RGB to 8-bit grayscale. Then, a threshold value was applied to represent the size of the lipid droplets most closely. This same threshold value was applied to all images. Finally, the percent area of the lipid droplets relative to the entire image was measured and the average value was reported for each animal. Samples were run in two batches. To account for batch-to-batch technical variation, values were normalized to the median value of the female samples for the strain in a specific batch. Groups were tested for significance by Wilcoxon rank sum test.

### Bulk RNA-seq Deconvolution

Bulk RNA-seq FASTQ files from Mckay *et al*. ^60^ were downloaded through NCBI SRA using NCBI SRA Toolkit (accession number GSE216369). FASTQ files were trimmed of adapters and low-quality reads were filtered using FASTP version 0.23.2 with default parameters^89^. FASTQ files were mapped to the killifish reference genome used in this study using STAR version 2.7.0e with parameters “--outFilterMultimapNmax 200; -- outFilterIntronMotifs RemoveNoncanonicalUnannotated; --alignEndsProtrude 10 ConcordantPair; --outSAMtype BAM SortedByCoordinate” ^90^. BAM files were summarized using featureCounts v2.0.4 ^91^ with parameters “--countReadPairs; -O; -p”.

Deconvolution was done using R package CSCDRNA version 1.0.3 under R version 4.1.2 with parameters “min.p = 0.3; markers = NULL” ^61^. Cell proportions were tested for significance using a Wilcoxon test rank-sum test. Predicted cell proportions and Benjamini-Hochberg-adjusted p-values are reported in **Supplementary Table S8**.

## Code Availability

All scripts used to analyze this dataset are available on the Benayoun lab GitHub at https://github.com/BenayounLaboratory/Killifish_Cell_Atlas.

## Data Availability

The raw FASTQ files have been deposited to the Sequence Read Archive under accession PRJNA952805. The final annotated Seurat object is available on FigShare (doi 10.6084/m9.figshare.22766894). Raw microscopy pictures of liver Oil-red-O staining are available on Figshare (doi: 10.6084/m9.figshare.22768076).

## Acknowledgments

Some panels were made with BioRender.com. We thank Jomille Jerez and Isabel Ollerton for assistance with killifish husbandry. We thank Dr. Minhoo Kim, and Juan Bravo, for feedback on our manuscript. We thank Cassandra McGill for advice on ORO image analysis using ImageJ. We thank Dr. Dario Valenzano for kindly sharing the ZMZ1001 turquoise killifish strain with us. This work was supported by NIA T32 AG052374 Postdoctoral Training Grant fellowship to B.B.T, a SURF (Summer Undergraduate Research Fund) from the USC Dornsife College of Letters, Arts, and Sciences to R.B., a Glenn/AFAR Grant for Junior Faculty and NIGMS R35 GM142395 to B.A.B.

The authors acknowledge the Center for Advanced Research Computing (CARC) at the University of Southern California for providing computing resources that have contributed to the research results reported within this publication (https://carc.usc.edu).

## Author contributions

B.B.T. and B.A.B. designed the study. B.B.T. and A.A. performed fish husbandry, dissected killifish tissues and isolated cells. B.B.T. and B.A.B. constructed 10xGenomics libraries. B.B.T., A.X. and B.A.B. performed computational analyses. A.J.J.L. performed RNA extraction and RT-qPCR of bulk tissue RNA. A.A., R.B. and K.H. performed liver ORO analysis. B.B.T. and B.A.B. wrote the manuscript with input from all authors. All authors edited and commented on the manuscript.

## Competing Interests

We declare no conflicts of interest.

## Supplementary Material

### Legends to Supplementary Figures

**Supplementary Figure S1. Tissue atlas quality control and cell type differentiation analysis using RNA velocity**

**(A)** Representative flow cytometry data from a male spleen group, showing gating strategy for singlet rate estimation. First, cells were distinguished from debris when plotting FSC-A *vs* SSC-A by drawing a common cell gate (shown) around all samples. This sample was 90.94% cells, which was typical of this protocol (**Supplementary Table S1**). Second, we noted events falling on a linear path when plotting FSC-A *vs*. FSC-H as singlets. This sample contained 99.98% singlets, which is typical of most singlet rates (>95%) (**Supplementary Table S1**). **(B)** Table of samples generated and used in the study. The number of animals used to derive a sample per sex per tissue per cohort is highlighted. Asterisks indicate if the sample was treated with the Dead Cell Removal Kit. Samples that were not used due to low cell viability or library construction failure are highlighted in red (**Supplementary Table S1**). **(C)** *Vitellogenin-1-like* expression is enriched in the female liver. *Vitellogenin-1-like* expression pattern across the tissue atlas as visualized by UMAP (left), as a violin plot by cell type and sex (middle) and as a violin plot by tissue of origin and sex (right). Note that vitellogenin is expressed at high levels exclusively in female cholangiocytes, endothelial cells, hepatocytes, and efferocytosing hepatocytes and is restricted in expression to the female liver. **(D)** Heatmap of marker gene expression levels of marker genes per cell type. For readability across more and less abundant cell types, we randomly sampled 200 cells per cell type to plot this heatmap. Marker genes show distinct cell type specificity. Original NCBI transcript names are in **Supplementary Table S9**.

**Supplementary Figure S2. Median number of detected genes across cell types and cell type proportions by tissue**

**(A)** The number of unique genes per cell type per cohort per sex. For each cell type for each cohort and sex, the median “nFeature_RNA” value was extracted from the completed tissue atlas Seurat object and plotted. No global trend in unique genes per cell type are clear, consistent with data homogeneity. **(B)** Pie charts of proportions of cell types per tissue by sex. Cell type proportions are overall similar in either sex. The kidney contains with most diverse set of cell types, while the blood is almost entirely comprised of erythrocytes.

**Supplementary Figure S3. MDS plot of pseudobulked transcriptomes across all cell types and both sexes**

**(A)** MDS plot of all the cell types in the complete tissue atlas. Each point is a cell type of a cohort and the sex is denoted by point shape. Points cluster much more strongly by cell type than by sex.

**Supplementary Figure S4. Analysis of sex-dimorphic gene expression patterns across cell types and cell type-specific functional enrichments**

**(A)** Correlation heatmap of gene expression log_2_ fold change in male vs. female cells. Particular cell types tend to cluster together, such as erythrocytes and hepatocytes. **(B)** Heatmaps of top 3 most recurrently significant female- and male-biased genes across the entire tissue atlas by differential gene expression analysis. *Vitellogenin-1-like*, the yolk precursor protein, is highly sex-dimorphic and biased towards females. **(C)** *hpx* RT-qPCR results from liver, muscle, and spleen. *hpx* is significantly upregulated in all male tissues besides muscle, in which it is trending towards upregulation. Significance in Wilcoxon rank sum test. **(D)** Dotplots of top five enriched GO terms by cell type for each sex. Some cell types such as B-cells and erythrocyte progenitors had terms only enriched in females, while myeloid progenitors had terms only enriched in males. Females tend to have the terms “response to hormone” and “response to lipid” enriched. Cell types represented in the MDS analysis but absent in GO dotplots did not have any significantly enriched GO terms.

**Supplementary Figure S5. Machine-learning analysis of sex-specific marker genes**

**(A)** “Out-of-Bag” machine learning performance for GBM and RF models across the 10 training cross validation folds. GBM and RF had similar (good) performance. **(B)** Violin plots of expression for the top sex-predictive genes from the machine-learning models. Sex-predictive genes tend to be highly sex-dimorphic across multiple cell types. *LINE-R2*, the most male-biased transcript expressed ubiquitously, is highlighted in light blue. **(C)** RT-qPCR results of *LINE-R2* expression in liver, muscle, and spleen. *LINE-R2* is not significantly upregulated in either sex as assessed by RT-qPCR in any tissue. Significance in Wilcoxon rank sum test. Original FishTE-DB name in **Supplementary Table S9**

**Supplementary Figure S6. Deconvolution of cell type proportions in a bulk RNA-seq liver dataset**

**(A)** Pie charts depicting the cell proportions found in each of the groups from McKay *et al*. compared to the tissue atlas liver proportions segregated by sex (pie charts repeated from the liver data in **Supplementary Figure S2B).** Cell type proportions are consistent between the original atlas and the deconvoluted predictions groups, suggesting good performance of the algorithm using our reference atlas. **(B)** Boxplot of cell type abundance estimates from male and female livers from McKay *et al*. For increased power, both livers from fish fed *ad libitum* and calorically-restricted diets are considered, but distinctly labelled. For each cell type, the cell type proportions from each sex were contrasted using a Wilcoxon-Rank sum test with Benjamini-Hochberg p-adjustment for multiple testing. Both dietary groups were included. Significance values can be found in **Supplementary Table S8**. B-cells and macrophages are significantly enriched in females (red, bold). Females: pink, Males: blue, *ad libitum* diet: circle, calorically-restricted diet: triangles. **(C)** Boxplot of cell type estimates from male and female livers from fish fed *ad libitum* and calorically-restricted diets from McKay *et al*. The proportion of each cell type belonging to a dietary group of the same sex in the liver was contrasted between dietary groups of the same sex. A Wilcoxon rank sum test with Benjamini-Hochberg p-adjustment was used to test for significance. There were no significant effects of dietary restriction on cell type composition of the livers, although this might be the result of insufficient power. Significance values can be found in **Supplementary Table S8**. Females fed *ad libitum* diet: dark pink, Males fed *ad libitum* diet: dark blue, females fed calorically-restricted diet: light pink, males fed calorically-restricted diet: light blue.

### Inventory of Supplementary Tables

**Supplementary Table S1: Singlet and cell viability data**

(A) Representative singlet rates for blood cells and dissociated cells from kidney, liver and spleen tissues. (B) Cell viability measurements for all cohorts. Samples excluded for low cell viability or library failure are emboldened and highlighted in red.

**Supplementary Table S2: Summary statistics for the full tissue atlas**

(A) Summary stats for the single-cell dataset

**Supplementary Table S3: Markers used for cell type annotation**

(A) Selected zebrafish cell type markers from Macaulay et al, Cell Rep, 2016. (B) Selected zebrafish cell type markers from Tang et al, JEM, 2017. (C) Selected zebrafish cell type markers from Athanasiadis et al, Nat Commun, 2017. (D) Selected zebrafish cell type markers from Carmona et al, Genome Res, 2017. (E) Selected zebrafish cell type markers from Farnsworth et al, Dev Biol, 2020 (F) Selected zebrafish cell type markers from Xia et al, PNAS, 2020. (G) Selected zebrafish cell type markers from Morrison et al, Hepatol Commun, 2022. (H) Selected zebrafish cell type markers from Wang et al, NAR, 2023. (I) Selected zebrafish cell type markers from Jiang et al, Front Cell Dev Biol, 2021. (J) Combined marker genes from Zebrafish cell atlases used for automated killifish atlas annotation.

**Supplementary Table S4: Differential gene expression results of pseudobulked cell type expression data**

(A) DESeq2 results of differential gene expression by sex using pseudobulked gene expression data from B-cells. (B) DESeq2 results of differential gene expression by sex using pseudobulked gene expression data from Endothelial Cells. (C) DESeq2 results of differential gene expression by sex using pseudobulked gene expression data from Erythrocyte Progenitors. (D) DESeq2 results of differential gene expression by sex using pseudobulked gene expression data from Erythrocyte. (E) DESeq2 results of differential gene expression by sex using pseudobulked gene expression data from Hepatocytes (Efferocytosing). (F) DESeq2 results of differential gene expression by sex using pseudobulked gene expression data from Hepatocytes. (G) DESeq2 results of differential gene expression by sex using pseudobulked gene expression data from Macrophages. (H) DESeq2 results of differential gene expression by sex using pseudobulked gene expression data from Mast Cells. (I) DESeq2 results of differential gene expression by sex using pseudobulked gene expression data from Multipotent Progenitors. (J) DESeq2 results of differential gene expression by sex using pseudobulked gene expression data from Myeloid Progenitors. (K) DESeq2 results of differential gene expression by sex using pseudobulked gene expression data from Neutrophils. (L) DESeq2 results of differential gene expression by sex using pseudobulked gene expression data from NK/T-cells. (M) DESeq2 results of differential gene expression by sex using pseudobulked gene expression data from Thrombocytes.

**Supplementary Table S5: GO results from pseudobulked cell type gene expression data**

(A) ClusterProfiler GSEA GOBP results using pseudobulked gene expression data from B-cells. (B) ClusterProfiler GSEA GOBP results using pseudobulked gene expression data from Endothelial Cells. (C) ClusterProfiler GSEA GOBP results using pseudobulked gene expression data from Erythrocyte Progenitors. (D) ClusterProfiler GSEA GOBP results using pseudobulked gene expression data from Hepatocytes Efferocytosing. (E) ClusterProfiler GSEA GOBP results using pseudobulked gene expression data from Hepatocytes. (F) ClusterProfiler GSEA GOBP results using pseudobulked gene expression data from Macrophages. (G) ClusterProfiler GSEA GOBP results using pseudobulked gene expression data from Myeloid Progenitors. (H) ClusterProfiler GSEA GOBP results using pseudobulked gene expression data from Neutrophils. (I) ClusterProfiler GSEA GOBP results using pseudobulked gene expression data from Thrombocytes.

**Supplementary Table S6: Machine learning feature importance**

(A) Feature importance summary from trained machine-learning models.

**Supplementary Table S7: RT-qPCR primers list**

(A) RT-qPCR primers used in this study.

**Supplementary Table S8: Liver deconvolution cell type proportions and statistical analyses**

(A) Results of liver deconvolution. Cell proportions for each NCBI deposited RNA-seq library are reported. (B) Wilcoxon test results comparing cell type proportions by sex from deconvoluted data from McKay et al. (C) Wilcoxon test results comparing cell type proportions by dietary group segregated by sex from deconvoluted data from McKay et al.

**Supplementary Table S9: NCBI and FishTEDB gene and TE name conversions**

(A) Killifish gene name conversion table from Figure 1, Supplementary Figure S1. (B) Killifish gene name conversion table from Figure 5, Supplementary Figure S5. (C) Killifish TE name conversion table from Figure 5, Supplementary Figure S5.

## Notes

### Competing Interest Statement

The authors have declared no competing interest.

