## Supplemental Figure S1 for "Widespread sex-dimorphism across single-cell transcriptomes of adult African turquoise killifish tissues"

A Representative singlet gating strategy

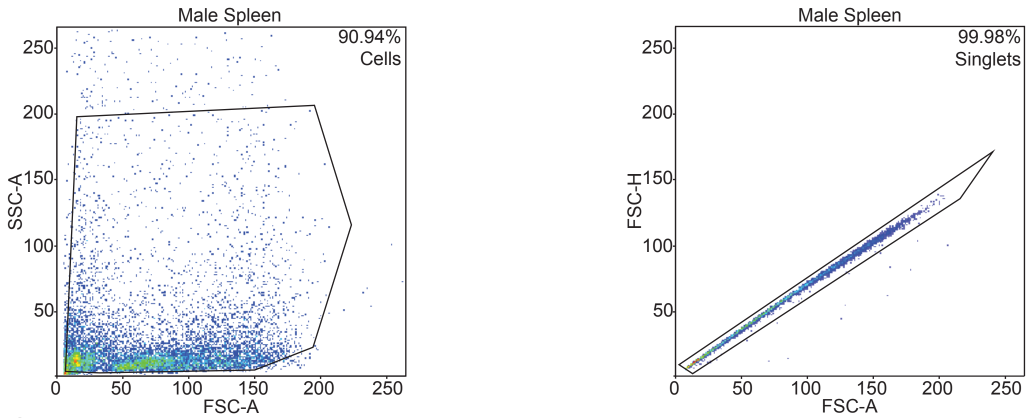


B Table of samples used in study

| | Cohort 1 | Cohort 2 | Cohort 3 |
|---|---|---|---|
| Blood | 4F | 3F | 3F* |
| | 3M | 3M | 3M* |
| Kidney | 4F | 3F | 3F* |
| | 3M | 3M | 3M* |
| Liver | 4F | 3F* | 3F* |
| | 3M | 3M* | 3M* |
| Spleen | 4F | 3F | 3F* |
| | 3M | 3M | 3M* |

*Dead cell removal kit used

C Vitellogen expression in female hepatocytes

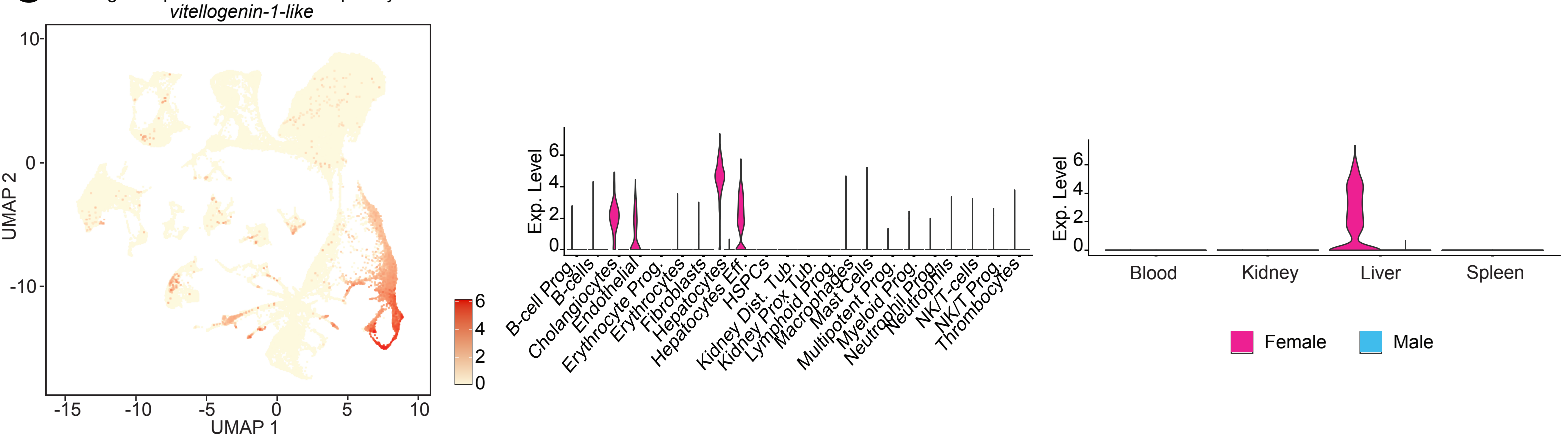


D Marker gene expression

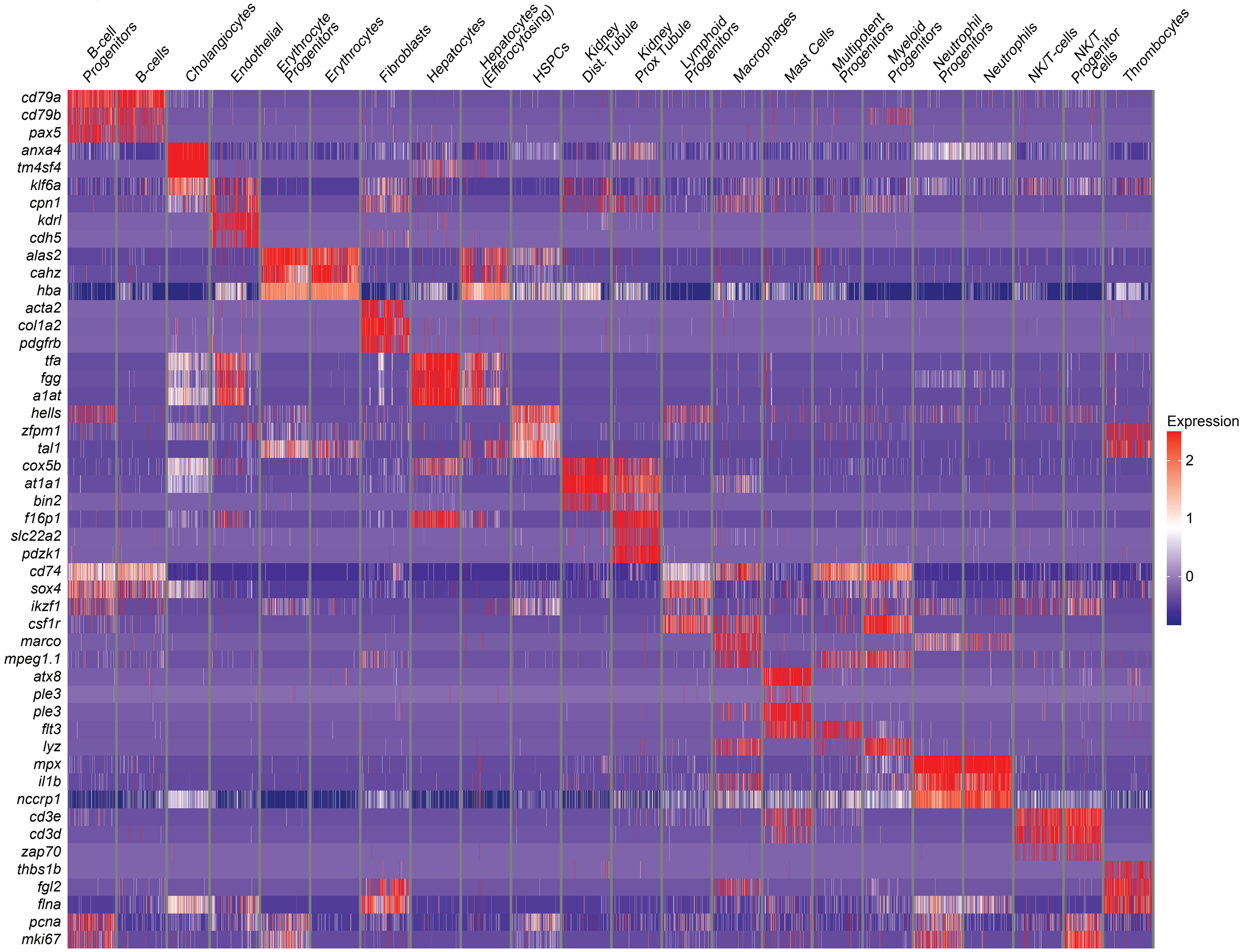
