## Supplementary figures and images for "Widespread sex-dimorphism across single-cell transcriptomes of adult African turquoise killifish tissues"

### Supplemental Figure S2

Figure S2

A Number of unique genes per cell type

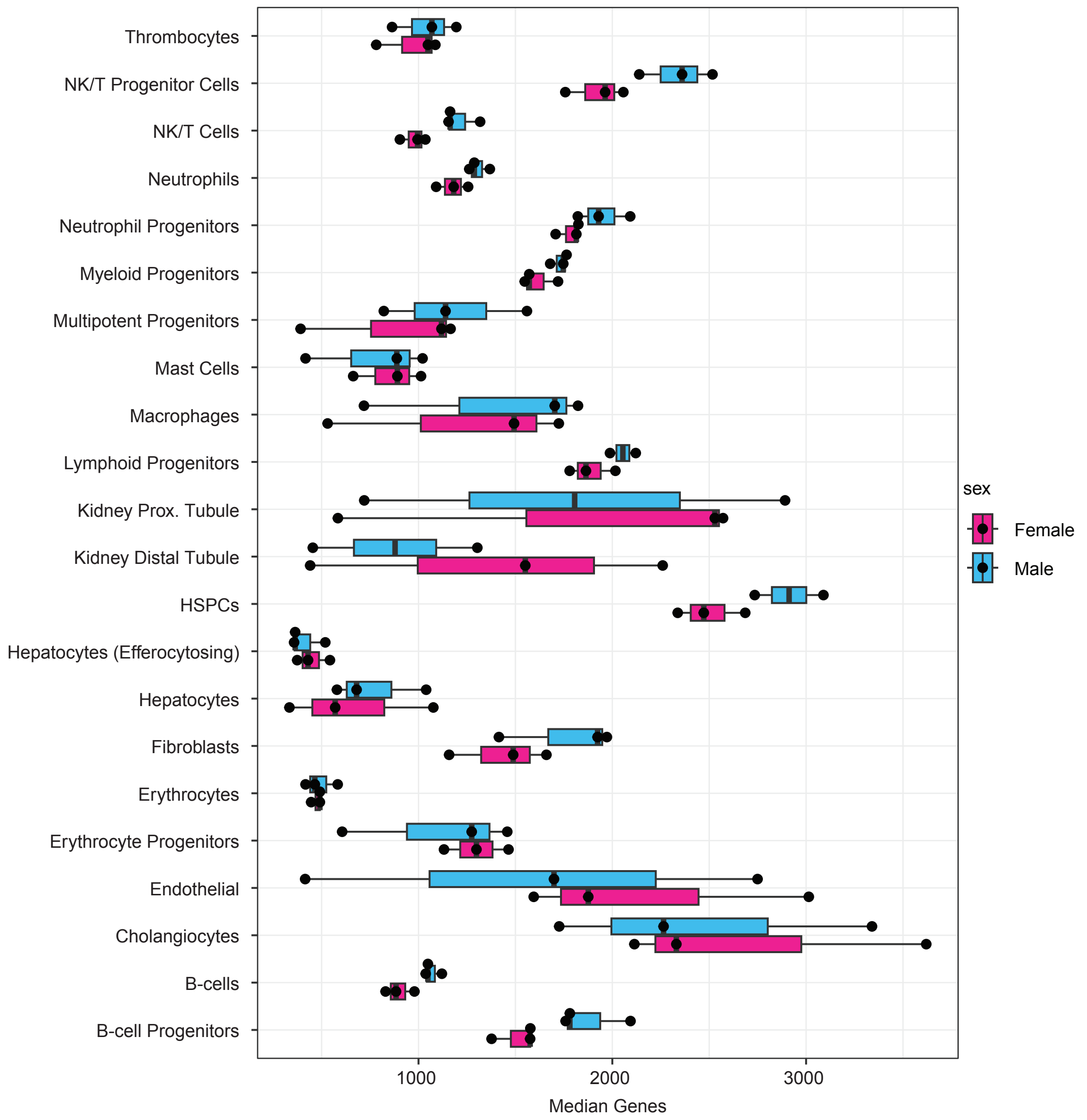


B Cell type proportions per tissue

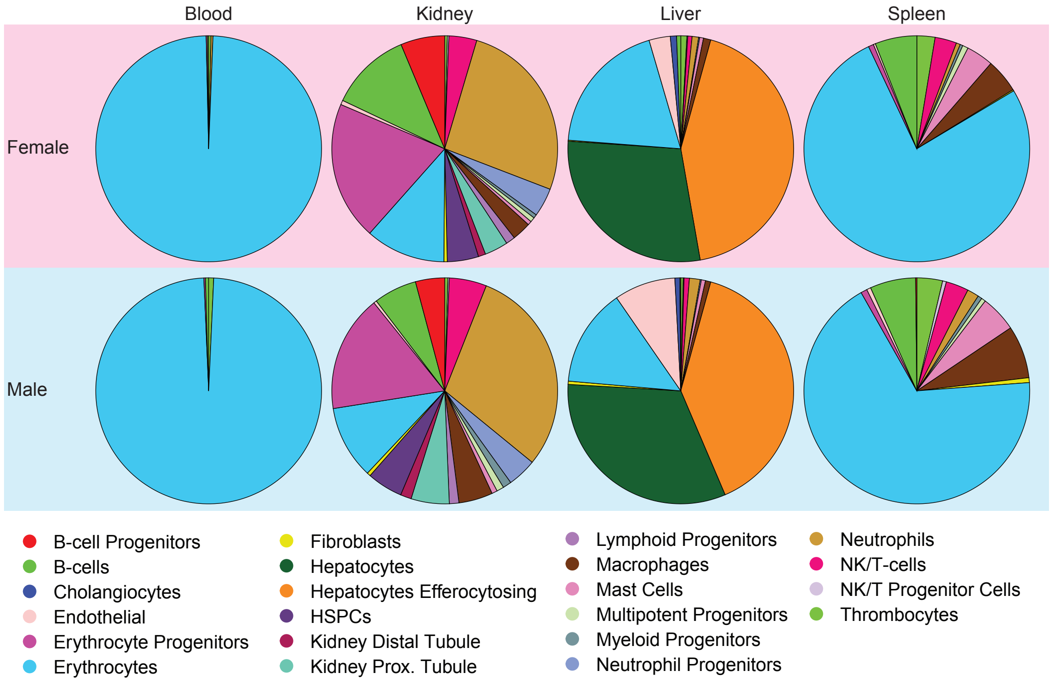

### Supplemental Figure S3

Figure S3

A MDS plot of killifish cell atlas

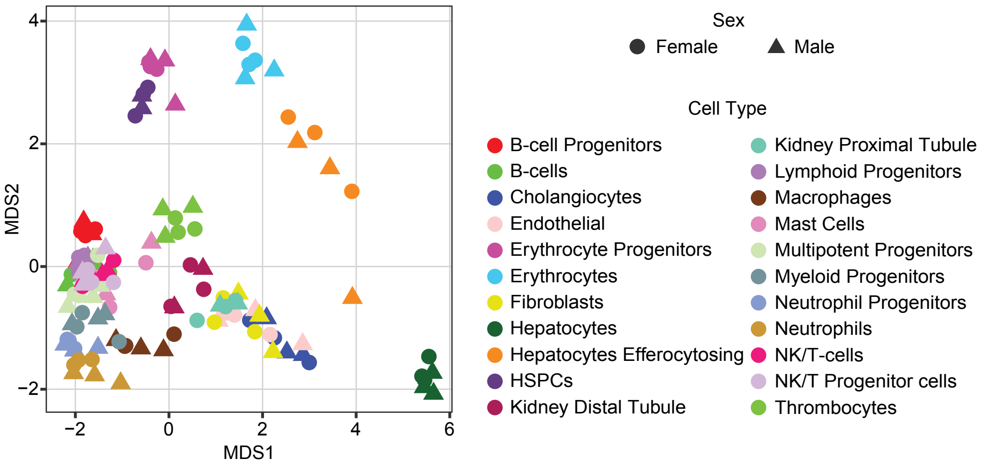

### Supplemental Figure S4

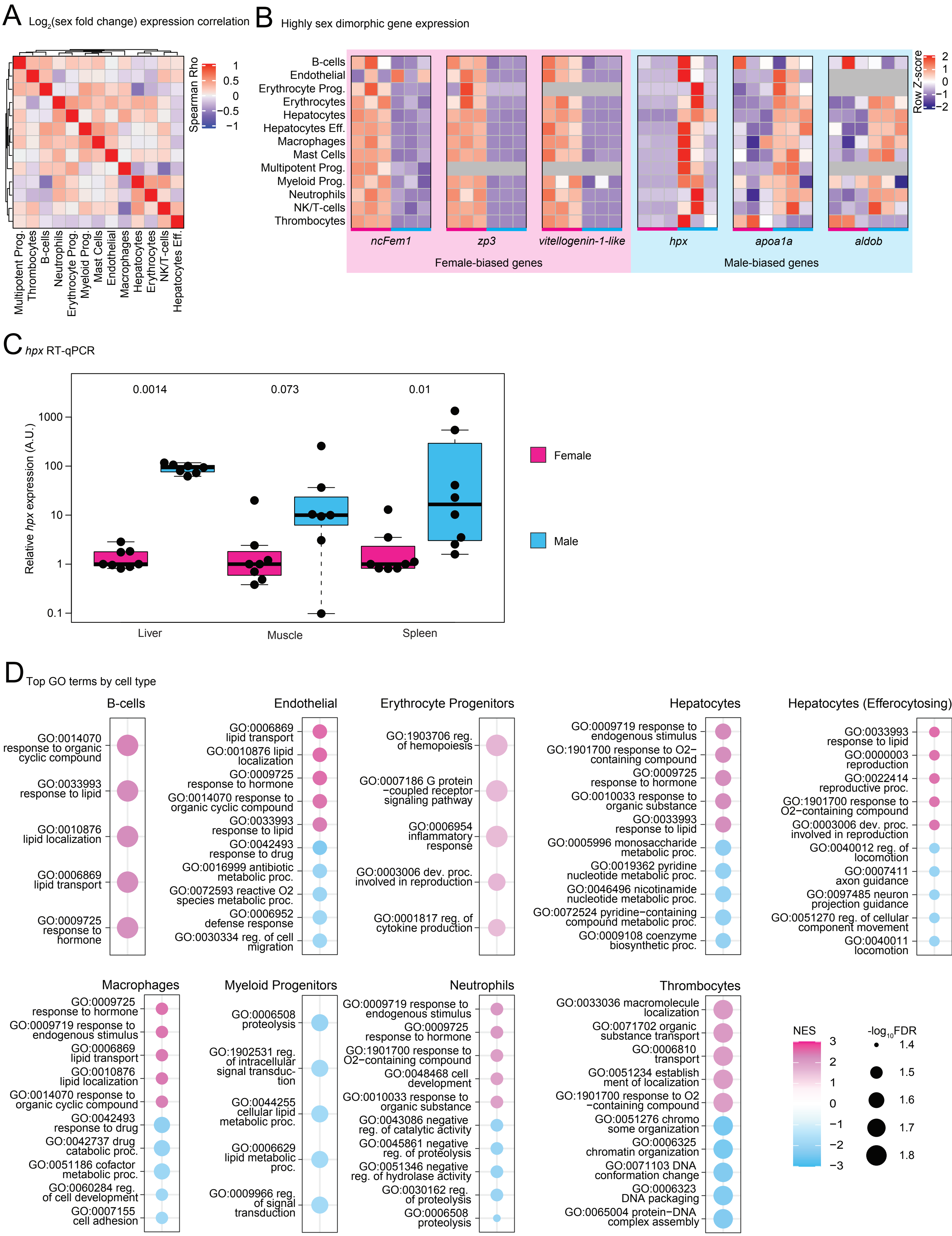

### Supplemental Figure S5

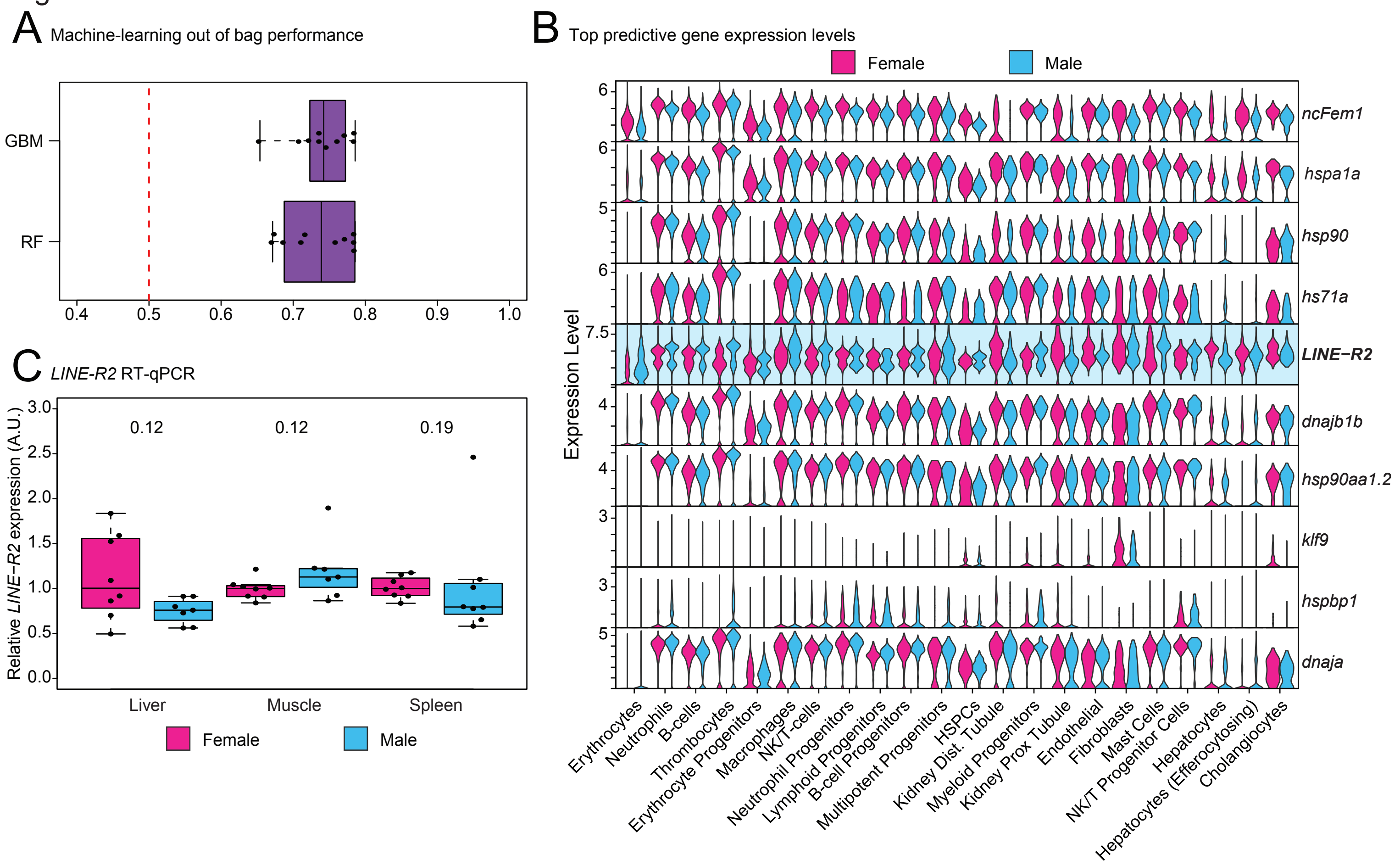

### Supplemental Figure S6

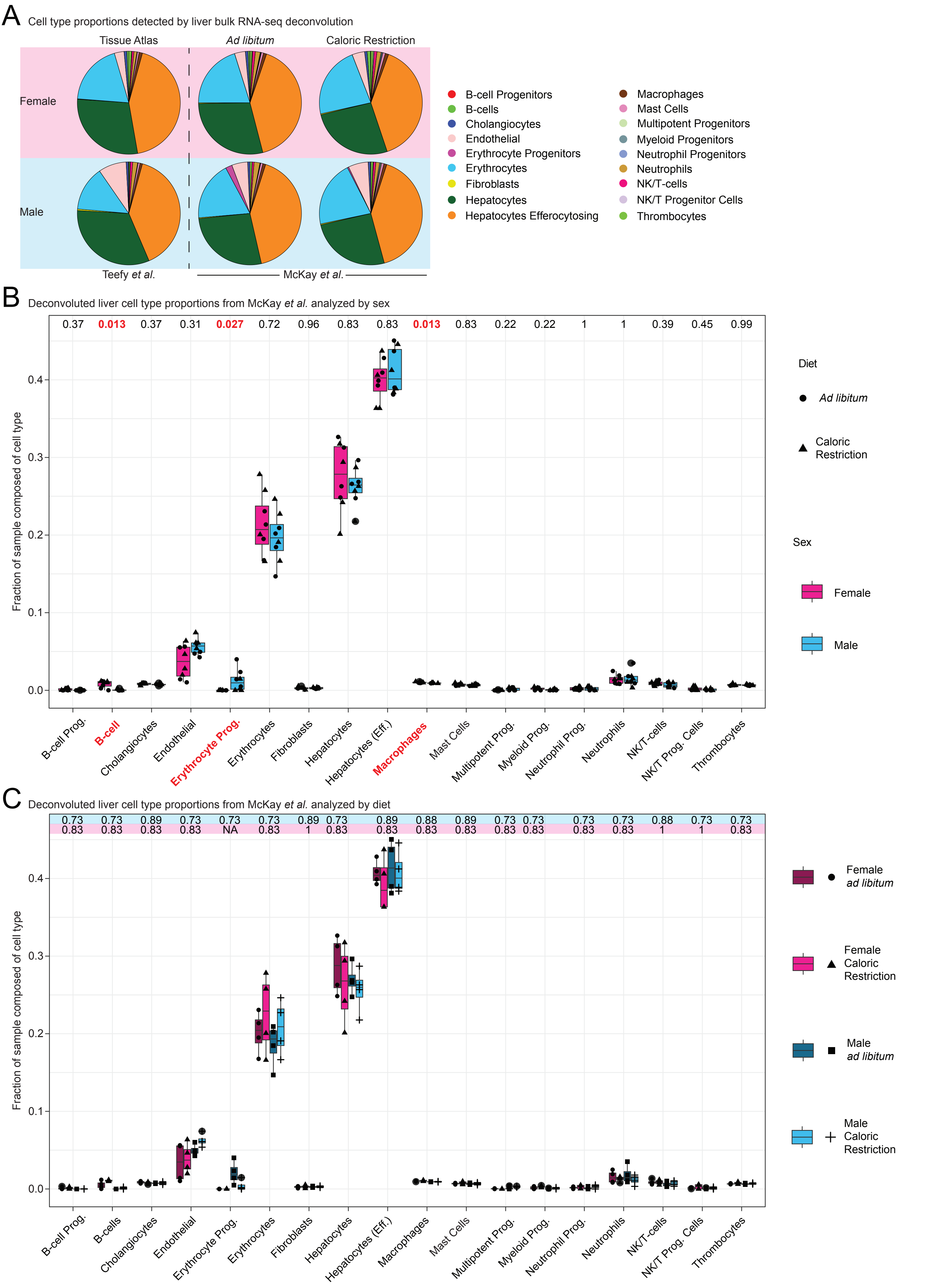
